# Granulins rescue inflammation, lysosome dysfunction, and neuropathology in a mouse model of progranulin deficiency

**DOI:** 10.1101/2023.04.17.536004

**Authors:** Jessica Root, Anarmaa Mendsaikhan, Srijita Nandy, Georgia Taylor, Minzheng Wang, Ludmilla Troiano Araujo, Paola Merino, Danny Ryu, Christopher Holler, Bonne M. Thompson, Giuseppe Astarita, Jean-François Blain, Thomas Kukar

## Abstract

Progranulin (PGRN) deficiency is linked to neurodegenerative diseases including frontotemporal dementia, Alzheimer’s disease, Parkinson’s disease, and neuronal ceroid lipofuscinosis. Proper PGRN levels are critical to maintain brain health and neuronal survival, however the function of PGRN is not well understood. PGRN is composed of 7.5 tandem repeat domains, called granulins, and is proteolytically processed into individual granulins inside the lysosome. The neuroprotective effects of full-length PGRN are well-documented, but the role of granulins is still unclear. Here we report, for the first time, that expression of single granulins is sufficient to rescue the full spectrum of disease pathology in mice with complete PGRN deficiency (*Grn^-/-^*).

Specifically, rAAV delivery of either human granulin-2 or granulin-4 to *Grn^-/-^* mouse brain ameliorates lysosome dysfunction, lipid dysregulation, microgliosis, and lipofuscinosis similar to full-length PGRN. These findings support the idea that individual granulins are the functional units of PGRN, likely mediate neuroprotection within the lysosome, and highlight their importance for developing therapeutics to treat FTD-*GRN* and other neurodegenerative diseases.

## INTRODUCTION

The granulin (*GRN)* gene encodes progranulin (PGRN), an ancient, evolutionarily conserved protein that is critical for brain health and neuronal survival.^1^ Specifically, haploinsufficiency of PGRN due to *GRN* mutations causes frontotemporal dementia (FTD), a common neurodegenerative disease in people under the age of 60.^2–4^ Complete deficiency of PGRN causes neuronal ceroid lipofuscinosis (NCL), a neurodegenerative lysosomal storage disorder (LSD).^5, 6^ Moreover, genetic variants in *GRN* decrease PGRN levels and have been associated with an increased risk of developing Alzheimer’s disease, Parkinson’s disease, or limbic-predominant age-related TDP-43 encephalopathy (LATE).^7–10^ Multiple therapeutic strategies are being pursued to treat the various neurodegenerative diseases associated with decreased PGRN.^11, 12^ Despite these advances, the fundamental function of PGRN is still unresolved and presents a roadblock for developing efficacious therapies for neurodegeneration.

PGRN is a ∼88 kDa secreted glycoprotein that is ubiquitously expressed and enriched in microglia and neurons in the brain.^13–15^ Mammalian PGRN is composed of 7.5 tandem repeat proteins, called granulins. Within PGRN, each granulin is joined together by short linear sequences or linkers, which can be cleaved by proteases to release mature granulins.^14, 16, 17^ We refer to each granulin numbered 1 through 7 based on the UniProtKB (P28799) database, rather than the colloquial A through G nomenclature. The relationship between the activity of PGRN and individual granulins has been debated and is still unclear. Multiple functions have been associated with full-length PGRN including cell growth, neurotrophic signaling, and anti-inflammatory activity. The pleiotropic activity of PGRN may occur through binding extracellular signaling receptors^4, 18^, however some PGRN-receptor interactions have not been widely replicated^19–21^, raising the possibility of other mechanisms of action. Furthermore, the function of individual granulins, also called epithelins, is also controversial and unresolved. Depending on the model system, the reported activity of granulins is paradoxical, ranging from enhancing neurotrophic activity^22^, to promoting inflammation^16^, inducing neurotoxicity^23^ or impairing lysosome function.^24^

The discovery by our lab, and others, that granulins are made constitutively inside lysosomes led us to reevaluate the functional relationship between PGRN and granulins.^25–27^ Because complete loss of granulins in humans and mice causes an LSD, with more severe neurodegeneration than observed in PGRN haploinsufficiency, we reasoned that granulins have an intra-lysosomal function. This idea is supported by the known function of other lysosomal proteins, such as saposins, which are generated from the prosaposin precursor protein.^28^ Here we test the hypothesis that PGRN serves as a precursor to granulins, which are the functional units that mediate lysosomal homeostasis and are neuroprotective. We used recombinant adeno-associated virus (rAAV2/1) to assess whether expression of individual granulins in the brain of PGRN deficient mice can correct disease-associated phenotypes. Our data show that neuronal expression of a single granulin fully rescues a range of phenotypes including lysosomal dysfunction, microglial activation, lipid abnormalities, and lipofuscin accumulation to the same extent as full-length PGRN. These findings provide compelling evidence that granulins are the bioactive subunits of PGRN and indicate that potential therapeutic approaches for FTD-*GRN* should consider their effect on granulin levels. Furthermore, this work supports the potential use of granulins themselves for the treatment of diseases associated with PGRN deficiency.

## RESULTS

### ICV injection of rAAV at birth leads to widespread expression of granulins, PGRN, and GFP throughout the mouse brain

To test the hypothesis that granulins are functionally active and neuroprotective, we utilized *Grn^-/-^* mice, which lack PGRN and develop pathology including neuroinflammation, lysosome dysfunction, and synaptic loss that increases with age. In these experiments, we compared human granulins 2 and 4, as previous studies suggested they have opposing functional activity.^29^ Furthermore, human granulin-2 (hGRN2) and human granulin-4 (hGRN4) share only 50% identity at the amino acid level, and we reasoned this would be sufficient to reveal differences in bioactivity if present (**Supp. Fig. 1A, B**). Human progranulin (hPGRN) and GFP served as positive and negative controls, respectively. For granulins and PGRN, we engineered expression constructs to include an N-terminal signal peptide (SP), to direct trafficking through the secretory pathway, followed by epitope tags (twin-Strep tag and V5 or FLAG) to facilitate detection preceding the coding region of interest (**Fig. 1A**). We validated these constructs in HeLa *GRN^-/-^* cells and found that hGRN2, hGRN4, and hPGRN were properly trafficked to the lysosome as well as secreted into the media (**Supp. Fig. 2**). Then, we generated recombinant hybrid Adeno-Associated Virus 2/1 (rAAV2/1) encoding hGRN2, hGRN4, hPGRN, and GFP and performed bilateral intracerebroventricular (ICV) injections of rAAV2/1 vectors into newly born (P0) litters of *Grn^-/-^* and *Grn^+/+^* mice (**Fig. 1B**). This experimental paradigm, termed somatic brain transgenesis (SBT), preferentially transduces neurons when using AAV vectors packaged in the capsid 1 serotype and leads to widespread and long-term expression of genes of interest in the mouse brain.^30–32^

**Figure 1.**
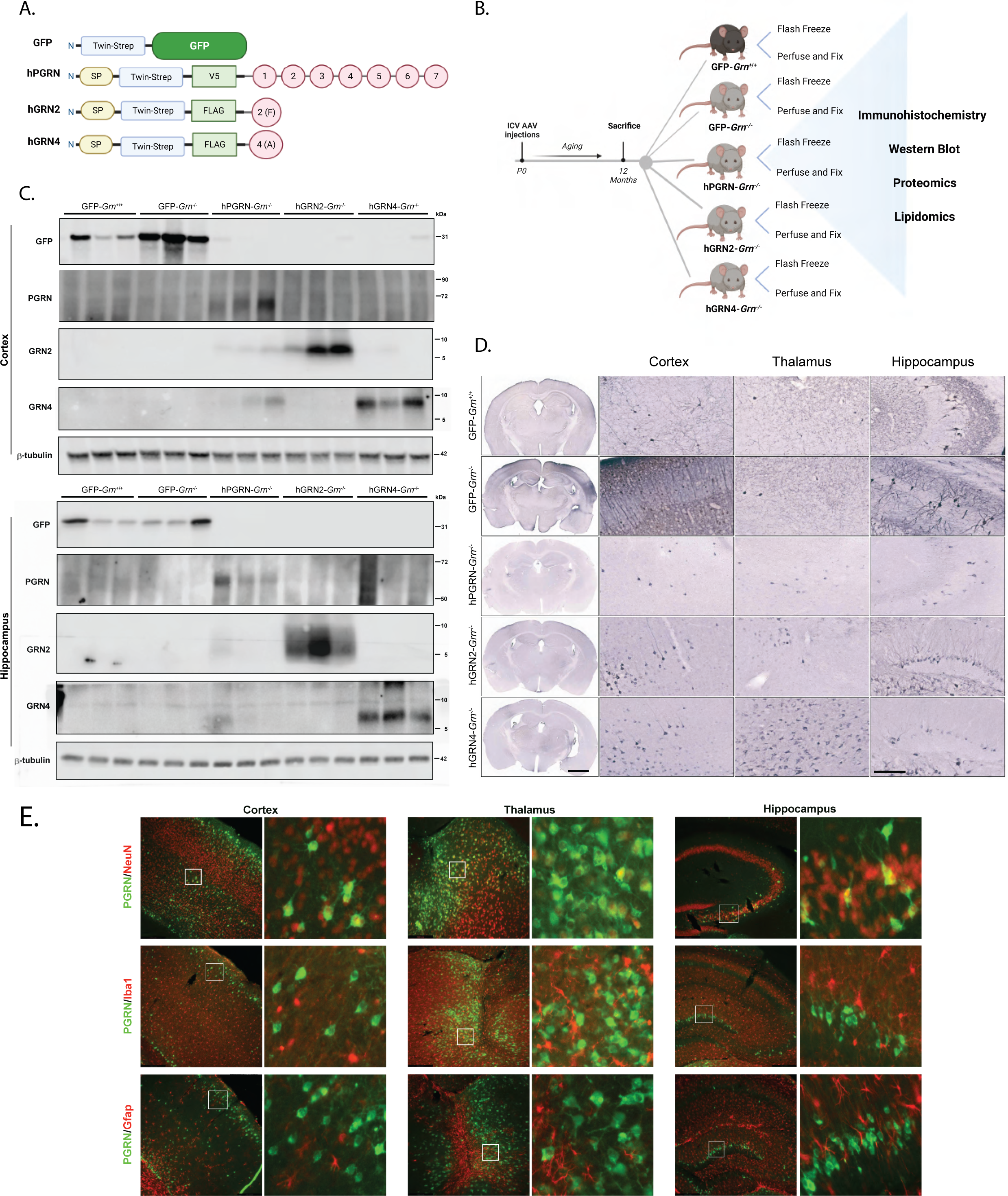
ICV injection of rAAV at birth leads to widespread expression of human granulins, PGRN, and GFP throughout the mouse brain. A) Diagram of all expression constructs including coding region of interest, domains, and epitope tags that were packaged into rAAV2/1 (twin-Strep-tag (TST); V5 epitope tag; FLAG epitope tag; SP= signal peptide; granulin-1; granulin-2; granulin-3; granulin-4; granulin-5; granulin-6; granulin-7). B) Diagram of experimental workflow including ICV injection of rAAV, mouse aging, sample collection, and sample analysis. C) Immunoblot verifying expression of encoded proteins following rAAV injection and aging. Cortical and hippocampal lysates were probed for GFP, hPGRN, hGRN2, hGRN4, with β-tubulin loading control. D) Representative immunohistochemistry (IHC) images for twin-Strep tag to visualize expression of GFP, hPGRN, hGRN2, and hGRN4 following AAV injection in whole coronal section plus magnified images of the cortex, hippocampus, and thalamus. Scale bar full coronal sections= 2 mm, regional zoom= 200 µm E) Representative immunofluorescent images co-staining for hPGRN and antibody markers for neurons (NeuN), microglia (Iba1), and astrocytes (GFAP) in the cortex, hippocampus, and thalamus of an hPGRN-*Grn^-/-^* mouse. White box highlights section of tissue enlarged on the right.

All mice were aged to 12-months, when substantial neuropathological changes are present in *Grn^-/-^* mice. Next, we characterized the distribution and expression of each rAAV2/1 vector throughout the brain of injected *Grn^-/-^* and *Grn^+/+^* mice. To confirm the specific identity of expressed proteins, we performed immunoblotting of lysates from flash frozen cortical and hippocampal brain tissue. Using specific antibodies, we confirmed expression of human progranulin in hPGRN-*Grn^-/-^* mice, human GRN2 in hGRN2-*Grn^-/-^* mice, human GRN4 in hGRN4-*Grn^-/-^* mice and GFP in GFP-*Grn^-/-^* and *Grn^+/+^* mice in both the hippocampus and cortex (**Fig. 1C**). Immunostaining of serial coronal sections for the twin-Strep tag, which is shared across expression constructs, visualized, and verified widespread expression of all encoded proteins in the hippocampus, thalamus, and cortex across injected mice (**Fig. 1D**).

Next, we assessed which cell types in the brain were transduced by rAAV2/1 and expressed specific transgenes. We utilized immunofluorescent staining to co-label hPGRN with the neuronal marker NeuN, the microglial marker Iba1, or the astrocytic marker GFAP in hPGRN-*Grn^-/-^*mice. We find that AAV-mediated transgene expression positively co-localized with the neuronal marker NeuN throughout the cortex, thalamus, and hippocampus. In contrast, we did not detect co-localization with Iba1 or GFAP (**Fig. 1E**). Thus, neonatal ICV injection of rAAV2/1 produced robust and stable neuronal expression of hGRN2, hGRN4, hPGRN, and GFP in mouse brains over the 12-month period of our experiments.

### Proteome-wide dysregulation in the thalamus of *Grn^-/-^* mice is ameliorated by expression of granulins

The thalamus is a major site of pathologic changes in *Grn^-/-^*mice^33^ and FTD-*GRN* patients^34, 35^, however the underlying pathogenic mechanisms are still poorly defined. To provide deeper insight into dysfunction of the thalamus caused by PGRN deficiency, we performed proteomics on flash frozen thalamic tissue of 12-month-old *Grn^-/-^* mice injected with rAAV2/1 encoding hGRN2, hGRN4, hPGRN, or GFP and *Grn^+/+^*mice injected with rAAV2/1 encoding GFP (**Fig. 1B**). Then, we performed quantitative proteomics of lysates of dissected thalamus using Tandem Mass Tagged (TMT) isobaric labeling followed by off-line electrostatic repulsion-hydrophilic interaction chromatography (ERLIC) fractionation prior to LC-MS/MS resulting in the identification and quantification of 9,255 proteins across all samples (**Supp. Fig. 3A**).

We next compared the GFP-*Grn^+/+^ and* GFP-*Grn^-/-^*thalamic proteomes to identify differentially expressed proteins. In GFP-*Grn^-/-^*mice we identified 131 proteins that increased and 9 proteins that decreased in abundance in the thalamus compared to GFP-*Grn^+/+^* mice (≥ 1.2-fold change; FDR q<0.05; **Fig. 2A**). Gene ontology (GO) analysis of the top 100 differentially expressed proteins using Metascape found a significant enrichment (−Llog10(p)L>L10) of proteins involved in lysosome function (Kegg mmu04142), glycosphingolipid metabolism (R-MMU-1660662), and protein catabolic processes in the vacuole (GO:0007039) (**Fig. 2B**). Some of the most significantly dysregulated proteins in *Grn^-/-^* mice included lysosomal hydrolases (Arsa, Gns, Hexa, Hexb, Manba) and proteases (Ctsd, Dpp7, Lgmn, Tpp1). Additionally, modules related to inflammatory processes were significantly enriched (−Llog10(p)L>L5), including MHC class II antigen presentation (R-MMU-2132295), regulation of complement cascade (R-MMU-977606), which include C1Qa, C1Qb, C1Qc, and C1Qb.

**Figure 2.**
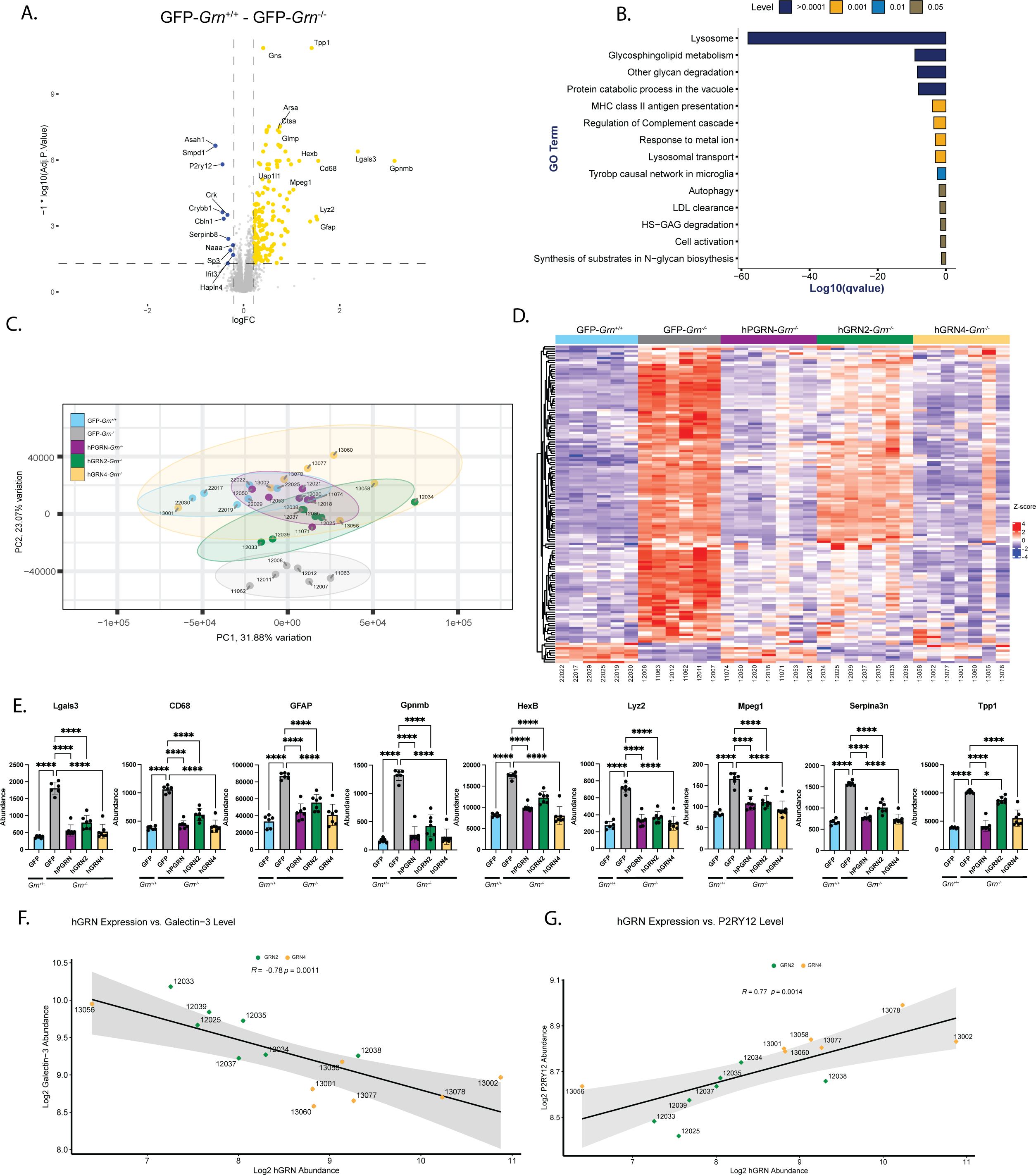
Granulins 2 and 4 prevent widespread protein dysregulation caused by PGRN deficiency in the thalamus of *Grn^-/-^* mice. A) Volcano plot of differentially expressed proteins in the thalamus of GFP-*Grn^-/-^* mice compared to GFP-*Grn^+/+^* mice. Upregulated protein in GFP-*Grn^-/-^* (yellow; right side) and downregulated in GFP-*Grn^-/-^* (blue; left side) are shown (FC>1.2, p<0.05). B) Bar graph of the most significantly enriched Gene Ontology (GO) terms describing the differentially expressed proteins in Fig. 2A (GFP-*Grn^-/-^* mice versus GFP-*Grn^+/+^* mice; FC 1.2 and adjusted p value 0.05). Displaying all significant changed modules (p value <0.05). C) Plot of largest principal components (PC1 vs. PC2) for GFP-*Grn^+/+^* mice (blue), GFP-*Grn^-/-^*mice (gray), hPGRN-*Grn^-/-^* mice (purple), hGRN2-*Grn^-/-^*mice (green), hGRN4-GFP-*Grn^-/-^* mice (yellow). Ellipses demarcate 95% confidence interval for assigned clusters. D) Heatmap of top 140 proteins (rows) differentially expressed between GFP-*Grn^-/-^* and GFP-*Grn^+/+^* across all treatment groups (columns). Quantification of individual proteins shown (Log_2_Z score transformed). Individual mouse ID numbers displayed below each column. E) Bar plots comparing correction of elevated levels of Lgals3, Cd68, Gfap, Gpnmb, Hexb, Lyz2, Mpeg1, Serpina3n, and Tpp1 in *Grn^-/-^* mice injected with GFP, hGRN2, hGRN4, or hPGRN. Data (protein abundance measured using TMT-based proteomics) mean ± SD. Significance was determined using a One-way ANOVA and corrected using Tukey’s post-hoc analysis. N=5-7 mice/group. * p < 0.05, ** p < 0.01, *** p < 0.001, **** p < 0.0001. (F, G) Correlation comparing hGRN2 and hGRN4 expression with Galectin-3 (F) (R=0.78 p=0.0011) or P2RY12 (G) (R=0.77, p=0.0014).

### Individual granulins rescue dysregulated proteins in the thalamus of *Grn^-/-^* mice

After characterizing differences in the proteome between *Grn^-/-^*and *Grn^+/+^* mice, we asked if expression of granulins or hPGRN could ameliorate changes observed in *Grn^-/-^* thalamus. First, principal component analysis (PCA) was performed, extracting 10 components from the proteomics dataset, accounting for 93% of variance (**Supp. Fig. 3B**). Comparing principal components 1 and 2 (PC1 and PC2) revealed a clear separation of GFP-*Grn^+/+^ and* GFP-*Grn^-/-^* samples with no overlap observed between groups (**Fig. 2C**). Samples from hGRN2-*Grn^-/-^*, hGRN4-*Grn^-/-^,* and hPGRN-*Grn^-/-^* mice overlap and cluster closer together with GFP-*Grn^+/+^* mice, revealing a shift towards wild-type mice, and away from *Grn^-/-^* mice, suggesting a general correction of altered protein levels.

To evaluate rescue of disease-linked phenotypes in more detail, we created a heatmap containing the 140 differentially expressed proteins from the GFP-*Grn^-/-^* and GFP-*Grn^+/+^* proteomics comparison and included hGRN2-*Grn^-/-^*, hGRN4-*Grn^-/-^*, hPGRN-*Grn^-/-^* samples (**Fig. 2D**). Visually the groups of *Grn^-/-^* mice treated with hGRN2, hGRN4, and PGRN are more like GFP-*Grn^+/+^* than GFP-*Grn^-/-^* mice. To provide a quantitative measurement of rescue, we compared expression levels of the most upregulated (2-fold; p< 0.001) proteins (GFAP, HEXB, SERPINA3N, TPP1, LYZ2, GPNMB, LGALS3, MPEG1, CD68) in the GFP-*Grn^-/-^* proteome across rAAV treatment groups. rAAV-mediated expression of either hGRN2, hGRN4, or hPGRN in *Grn^-/-^* mice significantly decreased expression levels back towards wild-type levels of all nine proteins, indicating correction of abnormally elevated proteins (**Fig. 2E**). This analysis provides strong evidence that expression of an individual granulin in the *Grn^-/-^* mouse brain can functionally substitute for the full length PGRN protein.

Of note, elevated proteins in *Grn^-/-^* mouse brains were not corrected as efficiently in hGRN2-injected groups compared to hGRN4 and hPGRN injected groups (e.g., Tpp1; **Fig. 2E**). One possible explanation for this result is that specific granulins are expressed at different levels between groups. To investigate this, we compared expression levels of hGRN2 and hGRN4 in the *Grn^-/-^* thalamic proteomics data set by examining a tryptic fragment of the twin Strep-FLAG tag shared between both proteins, revealing that hGRN4 expression was ∼2.5 fold higher than hGRN2 (**Supp. Fig. 3C**). We then asked whether the expression level of granulin-2 or granulin-4 correlated with phenotypic rescue. Notably, the abundance of hGRN2 and hGRN4 correlated with galectin-3 (R= -0.78; p=0.0011) (**Fig. 2F**) and P2RY12 (R= 0.77 p=0.0014) levels (**Fig. 2G**) in the *Grn^-/-^* thalamic proteome. Further, in individual mice higher levels of either hGRN2 or hGRN4 correlated with correction of altered protein levels, suggesting the decreased efficacy of hGRN2 is most likely due to lower expression levels and not function. Taken together, proteomic analysis of the thalamus *of Grn^-/-^* mouse reveals that rAAV-mediated expression of a single granulin ameliorates widespread protein dysregulation caused by loss of PGRN.

### Markers of Lysosomal Dysfunction are rescued by granulin expression across brain regions

To validate and extend the proteomics data, we analyzed tissue from additional, separate cohorts of rAAV2/1-injected *Grn^+/+^* and *Grn^-/-^* mice that were processed for immunohistochemistry, immunoblot (western blot), or lipidomics (**Fig. 1B**). Because “lysosome” was the most significant GO term in the GFP-*Grn^-/-^* thalamic proteome, we focused on two markers of lysosomal dysfunction, galectin-3 (LGALS3) and cathepsin Z (CatZ)^33^ (**Fig. 3A**).

**Figure 3.**
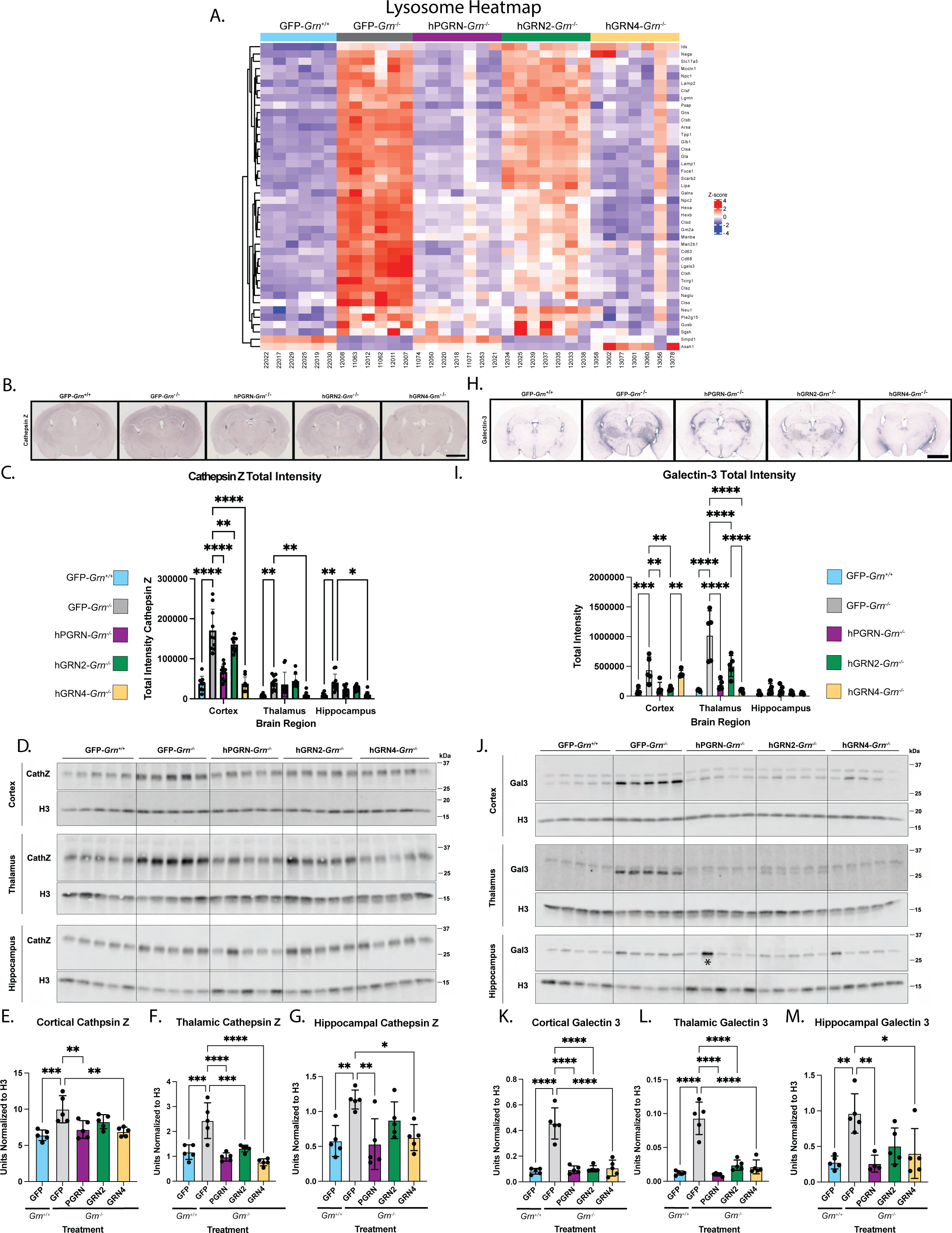
Individual granulins ameliorate dysregulated levels of lysosomal proteins, including cathepsin Z and galectin-3, in the brains of *Grn^-/-^*mice. A) Heatmap of differentially expressed (Log_2_Z score transformed) lysosomal proteins from GO module (Keeg mmu04142) in GFP-*Grn^-/-^* and GFP-*Grn^+/+^*mice. 42 proteins are included (rows) across mice from all treatment groups (columns). B) Representative images of cathepsin Z immunohistochemistry (IHC) staining of coronal sections of all rAAV injected groups (GFP, hPGRN, hGRN2, hGRN4). C) Quantification of cathepsin Z IHC signal in cortex, hippocampus, and thalamus. Data are mean±SD, significance determined with Two-Way ANOVA (two-way ANOVA _Region X Injection_ F (8, 133) = 17.37) followed by Tukey’s post-hoc analysis. N=5 mice/group. * p < 0.05, ** p < 0.01, *** p < 0.001, **** p < 0.0001 D) Immunoblot for cathepsin Z in cortical, hippocampal, and thalamic brain lysates from all injection groups. E) Quantification of immunoblot of cortical cathepsin Z normalized to H3. Data are mean±SD and significance determined by One-Way ANOVA (Tukey’s post-hoc analysis GFP-*Grn^-/-^* vs GFP-*Grn^+/+^* p=0.0009, GFP-*Grn^-/-^* vs hPGRN-*Grn^-/-^* p=0.0095, GFP-*Grn^-/-^* vs hGRN2-*Grn^-/-^* p=ns, GFP-*Grn^-/-^* vs hGRN4-*Grn^-/-^*p<0.0038). F) Quantification of immunoblot of thalamic cathepsin Z normalized to H3. Data are mean±SD and significance determined by One-Way ANOVA (Tukey’s post-hoc analysis GFP-*Grn^-/-^* vs GFP-*Grn^+/+^* p=0.0087, GFP-*Grn^-/-^* vs hPGRN-*Grn^-/-^* p=0.0046, GFP-*Grn^-/-^* vs hGRN2-*Grn^-/-^* p=ns, GFP-*Grn^-/-^* vs hGRN4-*Grn^-/-^*p<0.0175). G) Quantification of immunoblot of hippocampal cathepsin Z normalized to H3. Data are mean±SD and significance determined by One-Way ANOVA (Tukey’s post-hoc analysis GFP-*Grn^-/-^* vs GFP-*Grn^+/+^* p=0.0002, GFP-*Grn^-/-^* vs hPGRN-*Grn^-/-^* p<0.0001, GFP-*Grn^-/-^* vs hGRN2-*Grn^-/-^* p=0.0008, GFP-*Grn^-/-^* vs hGRN4-*Grn^-/-^*p<0.0001). H) Representative images of galectin-3 immunohistochemistry from coronal sections of all injection groups. I) Quantification of galectin-3 IHC signal in cortex, hippocampus, and thalamus. Data are mean±SD significance determined (two-way ANOVA _Region X Injection_ F (8, 60) = 11.95). followed by Tukey’s post-hoc analysis. N=5 mice/group. * p < 0.05, ** p < 0.01, *** p < 0.001, **** p < 0.0001. J) Immunoblot of galectin-3 in cortical, hippocampal, and thalamic brain lysates from all injection groups. K) Quantification of immunoblot of cortical galectin-3 normalized to H3. Data are mean±SD and significance determined by One-Way ANOVA (Tukey’s post-hoc analysis GFP-*Grn^-/-^* vs GFP-*Grn^+/+^*p<0.0001, GFP-*Grn^-/-^* vs hPGRN-*Grn^-/-^* p<0.0001, GFP-*Grn^-/-^* vs hGRN2-*Grn^-/-^* p<0.0001, GFP-*Grn^-/-^* vs hGRN4-*Grn^-/-^* p<0.0001). L) Quantification of immunoblot of thalamic galectin-3 signal normalized to H3. Data are mean±SD and significance determined by One-Way ANOVA (Tukey’s post-hoc analysis GFP-*Grn^-/-^* vs GFP-*Grn^+/+^* p<0.0001, GFP-*Grn^-/-^* vs hPGRN-*Grn^-/-^* p<0.0001, GFP-*Grn^-/-^* vs hGRN2-*Grn^-/-^* p<0.0001, GFP-*Grn^-/-^* vs hGRN4-*Grn^-/-^* p<0.0001). M) Quantification of immunoblot of hippocampal galectin-3 normalized to H3. Data are mean±SD and significance determined by One-Way ANOVA (Tukey’s post-hoc analysis GFP-*Grn^-/-^* vs GFP-*Grn^+/+^* p=0.002, GFP-*Grn^-/-^* vs hPGRN-*Grn^-/-^* p=0.0031, GFP-*Grn^-/-^* vs hGRN2-*Grn^-/-^* p=0.053, GFP-*Grn^-/-^* vs hGRN4-*Grn^-/-^*p=0.0132).

Cathepsin Z is a unique lysosomal cysteine protease that is upregulated in LSDs and neurodegenerative diseases.^36–38^ We performed IHC to examine the levels of cathepsin Z in hippocampal, thalamic, and cortical tissues of *Grn^-/-^* mice injected with rAAV2/1 expressing GFP, hGRN2, hGRN4, and hPGRN (n=5; one section/mouse) (**Fig. 3B**). Quantification of IHC staining in the cortex, thalamus, and hippocampus was performed using CellProfiler (**Supp. Fig 4)** and revealed a significant increase in cathepsin Z signal in GFP-*Grn^-/-^* mice across all regions examined (**Fig. 3C**). We found that the expression of hGRN2, hGRN4, and hPGRN corrected elevated levels of cathepsin Z in the cortex (**Fig. 3C**). In the thalamus and hippocampus only hGRN4 expression led to a statistically significant decrease in the level of cathepsin Z in these samples (**Fig. 3C**).

We then performed immunoblotting to provide a complementary measurement of cathepsin Z, in hippocampal, thalamic, and cortical tissue samples from separate cohorts (**Fig. 3D**). Cathepsin Z was increased in the GFP-*Grn^-/-^* mouse cortex, hippocampus, and thalamus compared to wild-type counterparts (**Fig. 3D, E-G**). In agreement with the proteomics analyses, cathepsin Z levels were normalized by expression of hGRN2, hGRN4, and hPGRN in the *Grn^-/-^* thalamus (**Fig. 3F**). Cathepsin Z levels were also decreased in the cortex and hippocampus by hGRN4 and hPGRN, while hGRN2 treatment trended lower, but did not reach significance (**Fig. 3E, 3G**).

We also assessed levels of galectin-3 (LGALS3), a beta-galactoside binding lectin that is recruited to damaged lysosomes^39^ to facilitate lysosomal repair.^40^ Immunostaining of 12-month-old GFP-*Grn^+/+^,* GFP-*Grn^-/-^,* hPGRN-*Grn^-/-^*, hGRN2*-Grn^-/-^*, and hGRN4-*Grn^-/-^* mouse coronal sections demonstrated that galectin-3 was increased in the thalamus and cortex of GFP-*Grn^-/-^* mice (**Fig. 3H**). Similarly, to cathepsin Z, expression of hGRN2, hGRN4, and hPGRN corrected elevated galectin-3 in the thalamus compared to GFP-*Grn^+/+^* mice (**Fig. 3I**).

These results were further validated by using immunoblot to measure galectin-3 levels in lysates of the cortex, thalamus, and hippocampus tissue from a separate cohort of rAAV2/1 injected mice (n=5) (**Fig. 3J**). We confirmed that galectin-3 was upregulated in cortical, thalamic, and hippocampus tissue lysates of GFP-*Grn^-/-^* mice compared to GFP-*Grn^+/+^* (**Fig. 3J, K-M**). Importantly, rAAV-mediated expression of hGRN2, hGRN4, and hPGRN reduced elevated galectin-3 expression in cortical and thalamic tissues (**Fig. 3K, L**). In hippocampal samples, hGRN4 and hPGRN significantly reduced elevated galectin-3 levels in *Grn^-/-^* mice (**Fig. 3M**). Together, these findings broaden the context of our proteomics data using IHC and immunoblot to confirm that expression of individual granulins reduce elevated levels of galectin-3 back to wild-type levels in the cortex and thalamus of 12-month-old GFP-*Grn^-/-^*mice.

In summary, we find that immunohistochemistry and immunoblot analysis confirm and extend our proteomics data, demonstrating that expression of hGRN2 or hGRN4 ameliorate elevated cathepsin Z and galectin-3 levels. These data provide additional evidence that a single granulin can functionally substitute for the activity of full-length PGRN.

### Microglial activation and inflammatory markers are reduced by hGRNs

While neuronal cell death is a hallmark of PGRN deficiency, the *GRN* gene is also highly expressed in microglia.^41^ Loss of PGRN in *Grn^-/-^*mice causes inflammation, astrocytosis, and microgliosis, which has been linked to synaptic loss and disease progression.^42, 43^ In the *Grn^-/-^* thalamic proteome, we find that many of the most dysregulated proteins are expressed by microglia including GPNMB, CD68, and P2RY12 (**Fig. 2A**). To examine this in more detail, we constructed a heatmap containing microglial activation markers found in the proteome of *Grn^+/+^ and Grn^-/-^* injected cohorts (**Fig. 4A**).^44, 45^ We find multiple markers of microglial activation including CD45 (PTPRC) and CD68 are upregulated in the GFP-*Grn^-/-^* thalamus (**Fig. 4** **B, D**). In addition, P2RY12, a marker of microglia homeostasis was down regulated in GFP-*Grn^-/-^* mice (**Fig. 4C**). Based on proteomics analyses, the elevation of CD45 was reduced by expression of hGRN2, hGRN4, and hPGRN, while depressed P2RY12 levels were increased by hGRN4 and hPGRN (**Fig. 4B, C**).

**Figure 4.**
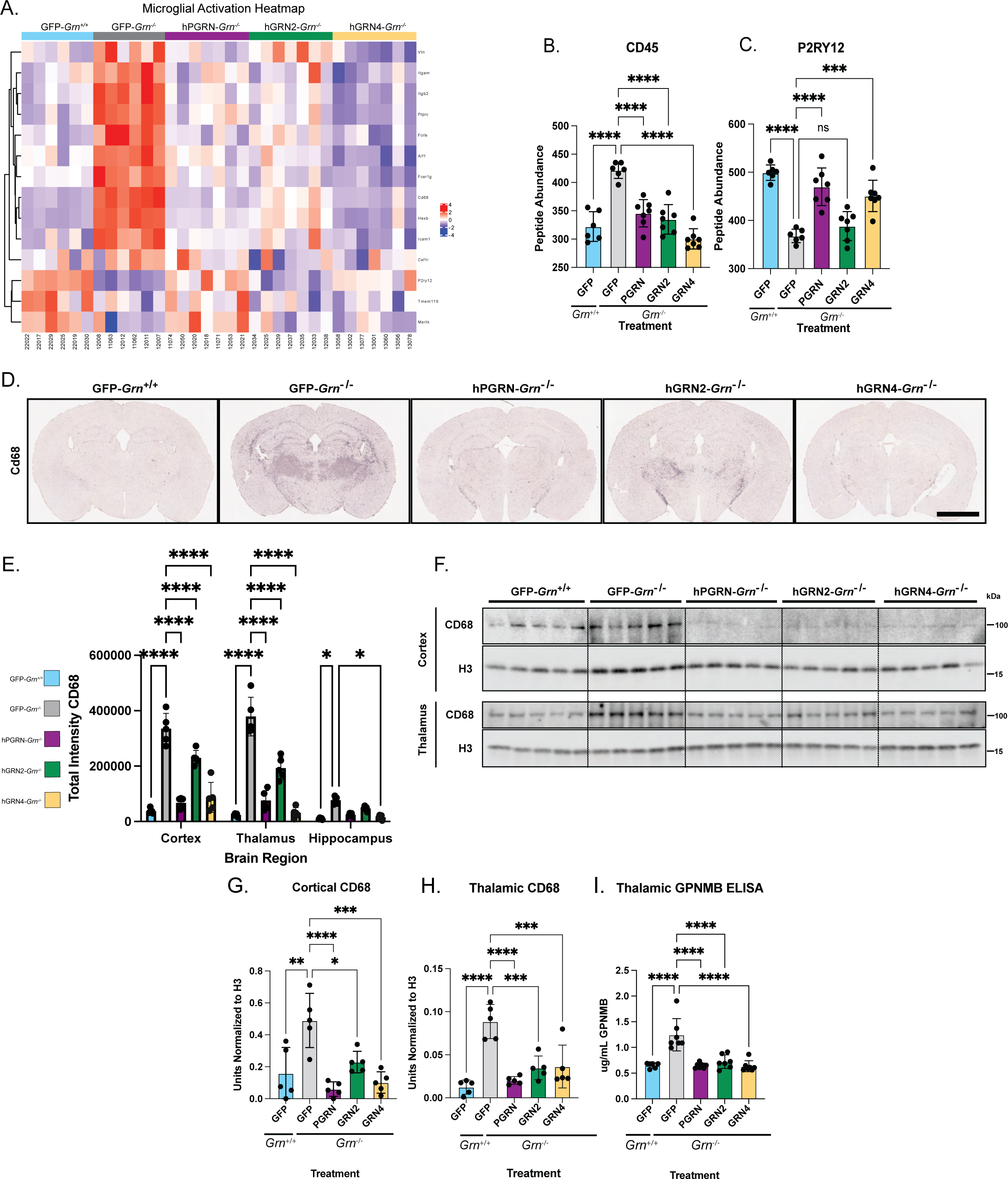
Granulins ameliorate multiple markers of microglial activation in *Grn^-/-^* mouse brains. A) Heatmap of differentially expressed (Log_2_Z score transformed) proteins associated with microglial activation and dysfunction^44, 45, 73^ (rows) in all treatment groups in GFP-*Grn^-/-^*compared to GFP-*Grn^+/+^* (columns). B) Proteomics abundance of CD45 (PTPRC) across all treatment groups (One-way ANOVA;Tukey’s post-hoc analysis GFP-*Grn^-/-^* vs GFP-*Grn^+/+^* p<0.0001, GFP-*Grn^-/-^* vs hPGRN-*Grn^-/-^* p<0.0001, GFP-*Grn^-/-^* vs hGRN2-*Grn^-/-^* p<0.0001, GFP-*Grn^-/-^* vs hGRN4-*Grn^-/-^*p<0.0001) C) Proteomics abundance of P2RY12 across all treatment groups. Data represented at mean ± SD One Way ANOVA (Tukey’s post-hoc analysis GFP-*Grn^-/-^* vs GFP-*Grn^+/+^* p<0.0001, GFP-*Grn^-/-^* vs hPGRN-*Grn^-/-^* p<0.0001, GFP-*Grn^-/-^* vs hGRN2-*Grn^-/-^* p= ns, GFP-*Grn^-/-^* vs hGRN4-*Grn^-/-^* p=0.0002). D) Representative images of CD68 immunohistochemistry of 12-month-old mouse coronal brain sections including all injection groups. E) Quantification of CD68 immunohistochemistry regions of interest cortex, hippocampus and thalamic signals quantified by CellProfiler. Data represented as mean±SD, significance was determined (Two Way ANOVA_Region X Injection_ F (8, 60) = 21.09). Followed by Tukey’s post-hoc analysis N=5 mice/group. * p < 0.05, ** p < 0.01, *** p < 0.001, **** p < 0.0001. F) Immunoblot of 12-month mouse cortical and thalamic brain tissue from all injection groups. G) Quantification of immunoblot of cortical CD68 signal normalized to H3. Data represented as mean±SD and significance determined by One-Way ANOVA (Tukey’s post-hoc analysis GFP-*Grn^-/-^* vs GFP-*Grn^+/+^*p=0.016, GFP-*Grn^-/-^* vs hPGRN-*Grn^-/-^* p<0.0001, GFP-*Grn^-/-^* vs hGRN2-*Grn^-/-^* p=0.0145, GFP-*Grn^-/-^* vs hGRN4-*Grn^-/-^*p=0.0003) H) Quantification of immunoblot of thalamic CD68 signal normalized to H3. Data represented as mean±SD and significance determined by One-Way ANOVA (Tukey’s post-hoc analysis GFP-*Grn^-/-^* vs GFP-*Grn^+/+^* p<0.0001, GFP-*Grn^-/-^* vs hPGRN-*Grn^-/-^* p<0.0001, GFP-*Grn^-/-^* vs hGRN2-*Grn^-/-^* p=0.0003, GFP-*Grn^-/-^* vs hGRN4-*Grn^-/-^*p=0.0004) I) Quantification of GPNMB levels in thalamic lysates measured using ELISA. Quantified data are mean±SD. Significance determined by one-way ANOVA (Tukey’s post-hoc analysis GFP-*Grn^-/-^*vs GFP-*Grn^+/+^* p<0.0001, GFP-*Grn^-/-^* vs hPGRN-*Grn^-/-^* p<0.0001, GFP-*Grn^-/-^* vs hGRN2-*Grn^-/-^* p<0.0001, GFP-*Grn^-/-^* vs hGRN4-*Grn^-/-^* p<0.0001)

To expand our investigation to additional brain regions and validate proteomics, we examined expression levels of CD68 (CD68), a type I transmembrane glycoprotein commonly used as a microglial activation marker^46^, using immunohistochemistry, and immunoblotting in additional cohorts of mice. Immunohistochemical staining for CD68 revealed robust and significant increases in CD68 in the cortex, hippocampus, and thalamus of GFP-*Grn^-/-^* mice (**Fig. 4D, E**). Expression of hGRN4 in *Grn^-/-^* mice lowered CD68 reactivity in all regions, while expression of hGRN2 significantly reduced levels in the thalamus and cortex, but not the hippocampus (**Fig. 4E**). Immunoblot of tissue lysates verified CD68 levels were increased in the cortex and thalamus of GFP-*Grn^-/-^* mice compared to GFP-*Grn^+/+^* mice, but below detection in the hippocampus (n=5; **Fig. 4F**). Similar to immunohistochemical analysis, rAAV expression of both hGRN2 and hGRN4 corrected elevated CD68 levels relative to GFP-*Grn*^-/-^ in the cortex (**Fig. 4G**) and in the thalamus (**Fig. 4H**).

Finally, we asked if expression of granulins corrected levels of glycoprotein non-metastatic melanoma protein B (GPNMB), the most elevated protein in the *Grn^-/-^* brain thalamic proteome (**Fig. 2A**), which was decreased by the expression of hGRN2 and hGRN4 in the thalamic proteomics (**Fig. 2E**). GPNMB is a type-1 transmembrane glycoprotein that we discovered to be highly upregulated by PGRN deficient microglia.^33^ The function of GPNMB in microglia is unknown, however GPNMB upregulation has been observed in activated damage-associated microglia^47^ and functionally linked to lysosomal stress and lipid accumulation.^48, 49^ We could not detect GPNMB via immunoblot, therefore, we quantified murine GPNMB levels in thalamic tissue lysates using a validated ELISA.^33^ Using this approach, we found that expression of either hGRN2 or hGRN4 corrected elevated GPNMB levels to the same extent as hPGRN in *Grn^-/-^* mice (**Fig. 4I**). In sum, proteomics and multiple orthogonal biochemical measurements reveal that expression of hGRN2, hGRN4, and hPGRN, especially in the thalamus, decrease microglial activation in *Grn^-/-^* mice.

### Lysosomal lipid dysregulation is rescued by a single granulin

The role of granulins in the lysosome is not fully understood. Previous studies identified lipid dysregulation in PGRN deficient animal models and FTD*-GRN* patient samples, suggesting lysosomal metabolism of lipids is impaired.^12, 50–52^ In particular, *Grn^-/-^* mice display decreased levels of bis(monoacylglycerol)phosphate (BMP), an atypical endo-lysosomal lipid, and increased levels of glucosylsphingosine (GlcSph), a substrate of glucocerebrosidase (GCase), which can be corrected by administration of exogenous full-length hPGRN.^53^ Additionally, ganglioside levels increase in brain tissues from *Grn^-/-^*mice and FTD-*GRN* patients ^54^, which may accumulate as secondary storage material, a phenomenon observed in many LSDs.^55, 56^ To validate these observations and determine whether expression of a single granulin or full-length hPGRN can correct them, we performed lipidomics and metabolomics analyses on the cortex of GFP-*Grn^+/+^*, GFP-*Grn^-/-^*, hPGRN-*Grn^-/-^*, hGRN2-*Grn^-/-^*, and hGrn4-*Grn^-/-^*mice.

These studies confirmed a decrease in the levels of BMP species and an increase in the levels of GlcSph species in GFP-*Grn^-/-^* mice compared with GFP-*Grn^+/+^* mice (**Fig. 5A**). In addition, we observed smaller, but significant increases of several gangliosides in the GFP-*Grn^-/-^* mouse brain (**Fig. 5A**), similar to what has been previously reported.^54^ AAV-mediated expression of hGRN4 and hPGRN in the *Grn^-/-^* mouse brain, but not hGRN2, significantly increased all measured BMP species back towards levels found in GFP-*Grn^+/+^* mice (**Fig. 5B**). Similarly, we found that both hGRN4 and hPGRN corrected the elevation of GlcSph (**Fig. 5C**) and gangliosides (**Fig. 5D**) in the GFP-*Grn^-/-^* mouse brain. hGRN2 showed a slight but non-significant reduction in GlcSph and ganglioside levels as well. It is unclear why hGRN2 did not rescue BMP and gangliosides to the same extent as hGRN4 and hPGRN. This could be due to lower expression of hGRN2 (**Supp. Fig. 3C**), or that different granulins may have specific functions in lysosomal lipid metabolism. Overall, these results demonstrate that a single granulin can correct multiple lysosomal lipids that are dysregulated in the *Grn^-/-^* mouse brain to the same extent as full-length hPGRN. Further research is required to understand the specific functions of different granulins on lysosomal lipid metabolism.

**Figure 5.**
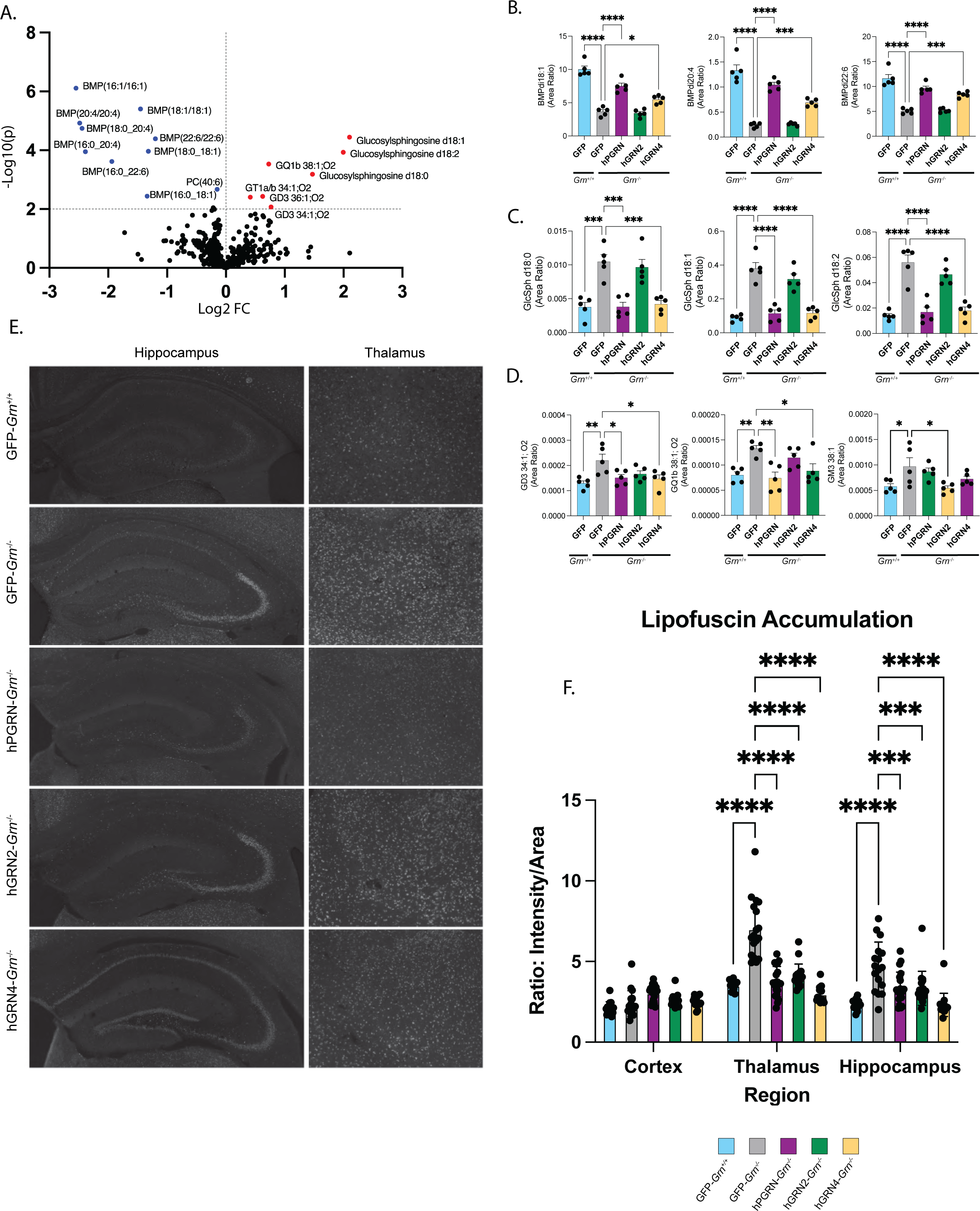
Granulins correct dysregulated lipids and prevent pathological accumulation of autofluorescent lipofuscin in *Grn^-/-^* mouse brains. A) Volcano plot of differential abundance of lipids and metabolites quantified in GFP-*Grn^-/-^* and GFP-*Grn^+/+^* mouse cortex. Lipids or metabolites upregulated in GFP-*Grn^-/-^* (red) and downregulated in GFP-*Grn^-/-^* (blue) are represented (p<0.1). B) Quantification of differentially abundant BMP species. Data represented as mean±SD, significance was determined by One-way ANOVA with Tukey’s post-hoc analysis. N=5-7 mice/group. * p < 0.05, ** p < 0.01, *** p < 0.001, **** p < 0.0001. C) Quantification of differentially abundant glycoshingosine species Data represented as mean±SD, significance was determined by One-way ANOVA with Tukey’s post-hoc analysis. N=5-7 mice/group. * p < 0.05, ** p < 0.01, *** p < 0.001, **** p < 0.0001. D) Quantification of differentially abundant gangliosides species. Data represented as mean±SD, significance was determined by One-way ANOVA with Tukey’s post-hoc ananlysis. N=5-7 mice/group. * p < 0.05, ** p < 0.01, *** p < 0.001, **** p < 0.0001 E) Representative images of lipofuscin autoflourescence as detected in the Cy5 channel from hippocampal and thalamic regions of coronal sections from all injection groups presented in grayscale. F) Quantification of fluorescent lipofuscin signal from cortex, hippocampus, and thalamus across all injected groups. Images were quantified in CellProfiler, and total intensity signal was divided by area of region assessed. Data represented as mean±SD significance determined with Two-Way ANOVA (two-way ANOVA_injectionXregion_ F _(8, 246)_ = 16.53 p<0.0001). Followed by Tukey’s post-hoc analysis N=5 mice/group. * p < 0.05, ** p < 0.01, *** p < 0.001, **** p < 0.0001

### Lipofuscin accumulation in *Grn^-/-^* brains is alleviated by expression of human granulins

Autofluorescent lipofuscin is a marker of lysosome dysfunction and is a neuropathologic feature of human FTD-*GRN* and *Grn^-/-^* mouse brain tissue.^57–59^ We set out to evaluate the extent and anatomical location of lipofuscin neuropathology in our rAAV-injected *Grn^-/-^*mouse brain cohorts. We imaged whole coronal sections using Cy5 excitation and emission filters to capture autofluorescence in GFP-*Grn^+/+^,* GFP-*Grn^-/-^,* hPGRN-*Grn^-/-^*, hGRN2*-Grn^-/-^*, and hGRN4-*Grn^-/-^* mice (n=5). Fluorescent signal in the cortex, hippocampus, and thalamic regions of all AAV injected mice was then quantified using CellProfiler (**Fig. 5E**). We observed a robust increase in lipofuscin in GFP-*Grn^-/-^* animals compared with GFP*-Grn^+/+^*in the thalamus and hippocampal regions, but not the cortex (**Fig. 5** **E, F**). Next, we found that expression of hGRN2, hGRN4, and hPGRN all decreased lipofuscin accumulation in the thalamus and hippocampus compared to GFP-*Grn^-/-^* mice (**Fig. 5** **E, F**). These data agree with our proteomic and lipidomic analyses and provide additional evidence that individual granulins can ameliorate widespread lysosomal dysregulation in *Grn^-/-^*mice including accumulation of lipofuscin, which has been linked to neurotoxicity and neurodegeneration in LSDs.^59^ This is the first report that a single granulin confers such broad beneficial effects in *Grn^-/-^* mice and provides compelling evidence that individual granulins are bioactive and neuroprotective.

## DISCUSSION

In this study, we find that delivery of an individual granulin using rAAV is equally efficacious as full-length hPGRN, correcting a variety of disease-linked neuropathology in *Grn^-/-^* mice. Proteomic analyses of the *Grn^-/-^* mouse thalamus at 12-months revealed wide-spread dysregulation of lysosomal hydrolases, lysosomal lipid metabolism, neuroinflammation, and proteostasis pathways, which can all be corrected by adding back hPGRN (∼88 kDa) or a single granulin (∼6k Da) subunit. This fills a critical gap in our knowledge of PGRN biology, strongly supporting the idea that individual granulins are the bioactive, functional components of PGRN. Moreover, our data suggest protein replacement with a single granulin may be a viable therapeutic approach in FTD caused by *GRN* mutations, which should be explored further.

Although it is well established that pathogenic *GRN* mutations decrease PGRN levels and ultimately cause neurodegeneration, the precise function of PGRN itself is still unclear. In general, the full-length PGRN protein has been thought to be directly neurotrophic, growth promoting, and anti-inflammatory. PGRN has been proposed to mediate these activities through binding and activation of extracellular signaling receptors^4, 18^, however this concept does not explain why complete lack of PGRN causes lysosome dysfunction and manifests as an LSD. Based on our discovery that PGRN is rapidly processed into individual granulins in the lysosome^60^, we tested the idea that the cleaved granulins themselves are active.

This is an important conceptual advance, because, prior to this study, granulins were thought to have the opposite activity of PGRN, potentially promoting inflammation^16^ and neurotoxicity^61^ while impairing lysosomal function^24^. In contradiction to these hypotheses, we find that long-term expression of two different granulins in *Grn^-/-^*mouse brain reduced multiple markers of neuroinflammation and glial activation, ameliorated the accumulation of lipofuscin, and broadly corrected dysregulated lysosomal proteins and lipids. Additional substantiation of our data comes from a study which found that PTV:PGRN, a brain penetrant form of hPGRN, is efficacious for multiple weeks after dosing *Grn^-/-^* mice, when PGRN has been completely cleared, suggesting that granulins made in the lysosome are stable and mediate prolonged efficacy^53^. Taken together, these findings suggest granulins have a central role regulating lysosomal function, lysosomal lipid metabolism, and may hold therapeutic potential for multiple neurodegenerative diseases associated with lysosomal dysfunction.^62–64^

A limitation of our study is that we only examined the efficacy of two of the seven granulins in *Grn^-/-^* mouse brains. We focused on hGRN2 and hGRN4 because they are 50% dissimilar, reported to have opposite function *in vitro*, and we were able to generate specific antibodies to assist experimental analysis. Considering this limitation, we found that rAAV-mediated expression of either hGRN2 or hGRN4 equally corrected major markers of lysosome dysfunction (galecin-3), microglial activation (Cd68, Gpnmb), and lipofuscin pathology in *Grn^-/-^*mouse brains. However, in some cases, such as correction of BMP and GlcSph lipids, hGRN2 was not as efficacious as hGRN4 or hPGRN. Proteomics quantification revealed that despite the injection of equivalent rAAV titers, hGRN2 was expressed at ∼2.5-fold lower levels than hGRN4, raising the possibility that insufficient expression of hGRN2 limited efficacy in the *Grn^-/-^* mouse brain. Alternatively, hGRN2 and hGRN4 may have different functions or binding partners in the lysosome explaining some of the observed differences. It is also possible that hGRN2 and hGRN4 may be differentially regulated or have different half-lives in the lysosome.^65^ Nevertheless, the shared ability of hGRN2 and hGRN4 to rescue many pathologic phenotypes in *Grn^-/-^* mice, strongly supports further investigation of the bioactivity of all granulins (1 to 7) *in vivo*.

The findings from our work have important implications for therapeutic development to treat FTD-*GRN* and other neurodegenerative diseases with PGRN deficiency. Multiple therapeutic strategies to increase PGRN levels in the CNS are being pursued for clinical development ranging from protein replacement^12^, to gene therapy^66^, and small molecule approaches.^67^ One approach aims to increase PGRN by depleting sortilin (*SORT1*), a PGRN lysosomal trafficking receptor, with an antibody (AL001) that has advanced to a phase 3 clinical trial (NCT04374136).^11^ Antibodies targeting the sortilin extracellular domain increase circulating levels of PGRN in mice^68^ but decrease lysosomal granulins in iPSC-derived neurons.^69^

Our data raise a concern with this approach, because anti-sortilin antibodies likely raise extracellular PGRN by reducing trafficking to the lysosome, leading to decreased production of intracellular granulins, which we find are functional and prevent lysosome dysfunction caused by PGRN deficiency. On the other hand, PGRN can be trafficked to the lysosome through alternative receptor pathways by binding prosaposin^70, 71^, which may reduce this concern, however it is unclear whether this occurs in the CNS following anti-sortilin treatment.^72^ At a minimum, our data raise a cautionary note that both PGRN and intra-lysosomal granulin levels should be measured when evaluating pre-clinical therapeutic approaches to treat PGRN deficiency in humans.

In conclusion, we find that neuronal expression of a single granulin can functionally substitute for the full length PGRN protein and correct a wide spectrum of disease-like phenotypes in *Grn^-/-^* mice including lysosome dysfunction, gliosis, dysregulated metabolism of lipids (BMP, GlcSph and gangliosides) and accumulation of lipofuscin. These findings support the idea that PGRN serves as a precursor to bioactive granulins, which are made in the lysosome, and directly mediate lysosomal homeostasis and neuroprotection. Key questions remain including whether all granulins can equivalently rescue pathologic phenotypes caused by PGRN deficiency. Additionally, the precise molecular function of granulins inside the lysosome, or whether each granulin has a unique or overlapping activity has yet to be resolved. From a therapeutic perspective, due to their small size, granulins may have advantages for treating FTD-*GRN* by crossing the blood-barrier more readily than PGRN due to their small size, although this needs to be empirically tested. Furthermore, therapies that aim to raise PGRN levels need to consider the impact on granulin levels throughout the CNS. Finally, our data strongly suggest that focused attention on the function of granulins inside the endosomal-lysosomal pathway is necessary to understand how granulins mediate lysosomal protein and lipid homeostasis and prevent neurodegeneration in FTD-*GRN* and related neurodegenerative disorders.

## Supporting information

Supplemental Figures 1 through 4

Supplemental Tables TMT Proteomics

Supplemental Tables Lipidomics and Metabolomics

## ACKNOWLEDGMENTS

We thank all the members of the Kukar lab, the Emory Center for Neurodegenerative Disease (CND), and the Emory Neuroscience (NS) graduate program for their support and helpful comments throughout this research project. We thank Dr. Shawn Ferguson (Yale) for the generous gift of HeLa *GRN^-/-^* cells. We greatly appreciate the kind support of Dr. Yona Levites and Dr. Todd E. Golde producing recombinant AAV. We thank Dr. Nicholas Seyfried, Duc Duong, and the Emory Proteomics core for excellent analytical services and technical expertise. Research reported in this publication was supported in part by the Emory Integrated Proteomics shared resource of Winship Cancer Institute of Emory University and NIH/NCI under award number P30CA138292. The content is solely the responsibility of the authors and does not necessarily represent the official views of the National Institutes of Health. This work was supported by the National Institutes of Health (NIH) National Institute of Neurological Disorders and Stroke (NINDS) grant R01NS105971, NIH National Institute of Aging (NIA) RF1AG079318 grant, a New Vision Research Investigator Award, the Alzheimer’s Drug Discovery Foundation (ADDF) and the Association for Frontotemporal Degeneration (AFTD), the Bluefield Project to Cure Frontotemporal Dementia, the BrightFocus Foundation, a gift from Arkuda Therapeutics, and an Emory School of Medicine (SOM) Dean’s Imagine, Innovate, and Impact (I^3^) Wow! Research Award Investigator grant. J.R. was supported by an NIH NINDS F31 Ruth L. Kirschstein National Research Service Award (NRSA) fellowship grant (F31NS117129).

## AUTHOR CONTRIBUTIONS

Conceptualization: J.R., C.H., G.T., and T.K.; Methodology: J.R., A.M., S.N., L.T.A., P.M., D.R., M.W., and G.T.; Formal analysis and investigation: J.R., A.M., S.N., B.M.T., L.T.A., P.M., M.W., and G.T., G.A., J-F.B.; Writing—original draft preparation: J.R., T.K.; Writing—review and editing: J.R., A.M., S.N., L.T.A., C.H., M.W., G.T., G.A. and J-F.B.; Funding acquisition: J.R. and T.K.; Supervision: T.K. All authors read and approved the final manuscript.

## FIGURE TITLES AND LEGENDS

## SUPPLEMENTAL FIGURE TITLES AND LEGENDS

**Supplemental Figure 1: Granulin Peptide Alignments and PID**

A. ClustalW alignment of Human granulin sequences. Visualized used ggmsa R package. Amino acids are color coded by their chemical properties.

B. Percent Identity heatmap derived from PID values calculated using granulin alignment output from figure S1A. Visualized using the Bio3D R package.

**Supplemental Figure 2: Validation that hPGRN, hGRN2, and hGRN4 are properly trafficked to the lysosome and secreted.**

A) Immunoblot of cell lysate and conditioned media collected from HeLa wild-type or *Grn^-/-^* cells expressing hPGRN, hGRN2, or hGRN4. Probed for hPGRN, hGRN2 and hGRN4.

**Supplemental Figure 3: Thalamic Proteomics**

A) Diagram of the proteomic thalamic workflow, displaying the number of proteins detected 9,255.

B) Assessment of Horn’s Parallel Analysis to determine how many components from the PCA to retain in downstream consideration.

C) Welch’s T-test comparing the abundance of granulin peptide detected in hGRN2-*Grn^-/-^* and hGRN4-*Grn^-/-^* mice thalamus (p-value=0.091) mean hGRN2= 288.4 mean hGRN4= 749.3

**Supplemental Figure 4: Cell Profiler Workflow**

A) Example of cropped ROIs from whole coronal section images. Bilateral images were collected from each section for each region.

B) Overview of CellProfiler workflow and output. Signal Area represents the regions quantified by pipeline.

## STAR Methods

### EXPERIMENTAL MODEL AND SUBJECT DETAILS

#### Mouse Model and Neonatal rAAV injections

The *Grn^−/−^* mice used in this study were purchased from the Jackson Laboratory (B6(Cg)-Grntm1.1Aidi/, IMSR Cat# JAX:013175, RRID:IMSR_JAX:013175) and generated as previously described. Mice were bred and housed in the Department of Animal Resources at Emory University and all work was approved by the Institutional Animal Care and Use Committee (IACUC) and performed in accordance with the Guide for the Care and Use of Laboratory Animals of the National Institutes of Health. Postnatal day 0 (P0) mouse pups (*GRN^+/+^* or *GRN^-/-^*) were injected with rAAV vectors.^32^ Briefly, P0 pups were cryoanesthetized in a nest protected by aluminum foil placed on ice for 5 min. One microliter of rAAV was injected intracerebroventricularly (ICV) into both hemispheres using a 10 ml Hamilton syringe with a 30-gauge needle. The pups were then placed on a heating pad with their original nesting material for 3–5 min and returned to their mother for further recovery. Mice were not sexed before injection, males and females were injected.

### METHODS

#### Production of recombinant adeno-associated virus

Four purified recombinant adeno-associated virus vectors (rAAVs) for injection were produced by plasmid transfection with helper plasmids in HEK293T cells. Briefly, the coding sequence of twin-Strep-GFP (GFP), twin-Strep-V5 human progranulin (hPGRN), twin-Strep-FLAG-granulin-2 with linker region 3 (hGRN2) and twin-Strep-FLAG-granulin-4 with linker region 5 (hGRN4) were subcloned from a pcDNA3.1 expression plasmid into pAAV. hPGRN, hGRN2, and hGRN4 all contain the native hPGRN signal peptide at the N-terminus. The AAV vectors express hPGRN, hGRN2, hGRN4, or GFP under the control of the cytomegalovirus enhancer/chicken β-actin promoter, a woodchuck post-transcriptional regulatory element, and the bovine growth hormone, poly(A), and were generated by plasmid transfection with helper plasmids in HEK293T cells. Forty-eight hours after transfection, the cells were harvested and lysed in the presence of 0.5% sodium deoxycholate and 50 U/ml Benzonase (Sigma, St. Louis, MO) by freeze thawing, and the virus was isolated using a discontinuous iodixanol gradient and affinity purified on a HiTrap HQ column (Amersham Biosciences, Arlington Heights, IL). The genomic titer of each virus was determined by quantitative PCR.

#### Collection of Brain Tissue

Mice were sacrificed after 12 months and brains were processed in two downstream pathways. Brains from half of the individuals from each cohort were immediately dissected from the skull and frozen at -80C. Whole brains were later thawed on ice and cortical, hippocampal, and thalamic sections were bulk dissected from the brain and frozen immediately at -80C. Remaining animals were transcardially perfused using ice cold PBS then fixed in methanol-free 4% PFA before dissecting all brains and storing in 4% PFA for 24 hours before transferring samples to 30% sucrose, which was replaced at 24 and 48 hours. The final storage solution was 30% sucrose and 1% sodium azide. Fixed tissue was stored at 4C until it was prepared for sectioning. Brain sectioning was performed using a freezing microtome set to 40 µm. Brains were frozen in ground dry ice, then mounted with 30% sucrose onto the pre-frozen sectioning stage, where serial sections were collected from the entire brain and stored in 30% sucrose, 30% ethylene glycol and 1% sodium azide. Frozen hippocampal and cortical brain samples from 12-month-old *Grn*^+/+^ (n=27) and *Grn*^-/-^ (n=44) mice were allocated for further processing.

#### Thalamic Proteomics Sample Preparation

Each tissue sample was homogenized in 300 μL of 8 M urea/100 mM NaHPO_4_, pH 8.5 with HALT protease and phosphatase inhibitor cocktail (Pierce) using a Bullet Blender (Next Advance) according to manufacturer protocols. Briefly, tissue lysis was transferred to a 1.5 mL Rino tube (Next Advance) with 350 mg stainless steel beads (0.9–2 mm in diameter) and blended for 5-minute intervals, two times, at 4°C. Protein supernatants were sonicated (Sonic Dismembrator, Fisher Scientific) three times for 5 seconds, with 15 second intervals of rest, at 30% amplitude to disrupt nucleic acids, in 1.5 mL Eppendorf tubes. Protein concentration was determined by BCA method, and aliquots were frozen at −80°C. Protein homogenates (200 µg) were treated with 1 mM dithiothreitol (DTT) at 25°C for 30Lminutes, followed by 5 mM iodoacetimide (IAA) at 25°C for 30Lminutes in the dark. Proteins were digested with 1:25 (w/w) lysyl endopeptidase (Wako) at 25°C for overnight followed by another overnight digestion with 1:25 (w/w) trypsin (Pierce) at 25°C after dilution with 50 mM NH_4_HCO_3_ to a final concentration of 1 M urea. The resulting peptides were desalted on a Sep-Pak C18 column (Waters) and dried under vacuum. All samples were across 2 batches and labeled with an 18-plex Tandem Mass Tag (TMTPro) kit (ThermoFisher, Lot numbers: UK297033 and WI336758) according to manufacturer’s protocol. Each TMT batch was desalted with 60 mg HLB columns (Waters) and dried via speed vacuum (Labconco). Dried samples were re-suspended in high pH loading buffer (0.07% vol/vol NH_4_OH, 0.045% vol/vol FA, 2% vol/vol ACN) and loaded onto a Water’s BEH column (2.1 mm x 150 mm with 1.7 µm particles). A Vanquish UPLC system (ThermoFisher Scientific) was used to carry out the fractionation. Solvent A consisted of 0.0175% (vol/vol) NH_4_OH, 0.01125% (vol/vol) FA, and 2% (vol/vol) ACN; solvent B consisted of 0.0175% (vol/vol) NH_4_OH, 0.01125% (vol/vol) FA, and 90% (vol/vol) ACN. The sample elution was performed over a 25 min gradient with a flow rate of 0.6 mL/min with a gradient from 0 to 50% solvent B. A total of 192 individual equal volume fractions were collected across the gradient. Fractions were concatenated to 96 fractions and dried to completeness using vacuum centrifugation. Dried peptide fractions were resuspended in 20 μl of peptide loading buffer (0.1% formic acid, 0.03% trifluoroacetic acid, 1% acetonitrile). Peptide mixtures (2 μl) were separated on a self-packed C18 (Dr. Maisch) fused silica column (15 cm × 150 μm internal diameter) by a Dionex Ultimate rsLCnano and monitored on a Fusion Lumos mass spectrometer (ThermoFisher). Elution was performed over a 42 min gradient at a rate of 1250 nl/min with buffer B ranging from 1% to 99% (buffer A: 0.1% formic acid in water, buffer B: 0.1% formic in 80% acetonitrile). The mass spectrometer cycle was programmed to collect at the top speed for 3 s cycles. The MS scans (410-1600 m/z range, 400,000 AGC, 50 ms maximum ion time) were collected at a resolution of 60,000 at m/z 200 in profile mode. HCD MS/MS spectra (0.7 m/z isolation width, 35% collision energy, 125,000 AGC target, 86 ms maximum ion time) were collected in the Orbitrap at a resolution of 50000. Dynamic exclusion was set to exclude previous sequenced precursor ions for 20 s within a 10-ppm window. Precursor ions with +1 and +8 or higher charge states were excluded from sequencing.

#### Thalamic Proteomics Data Processing

All raw files were analyzed using the Proteome Discoverer Suite (v.2.4.1.15, ThermoFisher). MS/MS spectra were searched against the UniProtKB mouse proteome database (downloaded in August 2020 with 91417 total sequences) supplemented with 4 variant sequences (twin-Strep-GFP, hGRN2, hGRN4, and hPGRN). The Sequest HT search engine was used to search the RAW files, with search parameters specified as follows: fully tryptic specificity, maximum of two missed cleavages, minimum peptide length of six, fixed modifications for TMTPro tags on lysine residues and peptide N-termini (+304.207LDa) and carbamidomethylation of cysteine residues (+57.02146LDa), variable modifications for oxidation of methionine residues (+15.99492LDa), serine, threonine and tyrosine phosphorylation (+79.966LDa) and deamidation of asparagine and glutamine (+0.984LDa), precursor mass tolerance of 10Lppm and a fragment mass tolerance of 0.05LDa. Percolator was used to filter peptide spectral matches and peptides to an FDR <1%. Following spectral assignment, peptides were assembled into proteins and were further filtered based on the combined probabilities of their constituent peptides to a final FDR of 1%. Peptides were grouped into proteins following strict parsimony principles.

#### Differential expression analysis

Differentially enriched or depleted proteins (pL≤L0.05) were identified by one-way ANOVA with post-hoc Tukey HSD test comparing five groups: GFP-Grn^+/+^, GFP-*Grn^-/-^,* PGRN-*Grn^-/-^,* GRN2-*Grn^-/-^,* GRN4-*Grn^-/-^* mice. Differential expression of proteins was visualized with volcano plots generated using the ggplot2^74^ package in Microsoft R Open v3.4.2. Significantly differentially expressed proteins were determined by both having a pL≤L0.05 and a fold change difference of greater than log2(1.25) or less than −Llog2(1.20) (a minimum 1.2-fold change).

#### Proteomics Analysis and Visualization

Differential Expression data from comparisons GFP-Grn^+/+^ vs GFP-*Grn^-/-^,* GFP-*Grn^-/-^* vs hPGRN-*Grn^-/-^,* GFP-*Grn^-/-^* vs hGRN2-*Grn^-/-^,* and GFP-*Grn^-/-^,* versus hGRN4-*Grn^-/-^* including adjusted p values, and abundance values were imported into Quickomics, an R-shiny powered proteomics analysis and visualization tool.^75^ GIS internal standards were removed from the data set and Heatmaps were created filtering proteins from the GFP-Grn^+/+^ vs GFP-*Grn^-/-^* comparison with and adjusted p-value of <0.05 and a fold change value of at least 1.2 or 20%. Clustering was performed grouping proteins by the similarity across the sample ID using a k-means approach. Other visualizations created in Quickomics include 2-Way DEG plots and PCA visualizations. Additional PCA analysis was undertaken in R using the PCAtools package.^76^

#### Gene ontology (GO)

Genes IDs identified from proteins determined to be differentially abundant (adjusted p-value 0.05, FC 1.2) between GFP-Grn^+/+^, GFP-*Grn^-/-^*mice were input into the Metascape Gene Ontology Analysis tool (https://metascape.org).^77^ Express Analysis was conducted and the top 50 Ontology Terms were collected.

#### Lipidomics and Metabolomics

##### Sample preparation for lipidomics and metabolomics analyses

During tissue collection, the cortex was dissected, weighed, and flash frozen. Each frozen cortex was pulverized into a homogenous powder, and roughly 30 mg of each cortex powder sample was used to extract lipids. Methanol spiked with internal standards (see LCMS methods below) was added to each sample and homogenized with FastPrep-24™ 5G bead beating grinder and lysis system using Lysing Matrix D tubes with CoolPrep™ adapter (MP Biomedicals) for 40 seconds at a speed of 6 m/s. The methanol fraction was then isolated via centrifugation (20 minutes at 4°C, 14,000 x g), followed by transfer of supernatant to a 96 well plate. After a 1 h incubation atL20°C followed by an additional centrifugation (20 minutes, 4,000 x g at 4°C), methanol was transferred to glass vials for LCMS analysis.

##### Lipidomics analysis

Lipid analyses were performed by liquid chromatography on an ExionLC (Sciex) coupled with electrospray mass spectrometry TripleQuad 7500 (Sciex). For each analysis, 1 µL of the sample was injected on a Premier BEH C18 1.7 µm, 2.1×100 mm column (Waters) using a flow rate of 0.25 mL/min at 55°C. For positive ionization mode, mobile phase A consisted of 60/40 (vol/vol) acetonitrile/water with 10 mM ammoniumLformateL+ 0.1% formic acid; mobile phase B consisted of 90/10L (vol/vol) isopropyl alcohol/acetonitrile with 10 mM ammoniumLformateL+ 0.1% formic acid. For negative ionization mode, mobile phase A consisted of 60/40 (vol/vol) acetonitrile/water with 10 mM ammonium acetate; mobile phase B consisted of 90/10L(vol/vol) isopropyl alcohol/acetonitrile with 10 mM ammonium acetate. The gradient was programmed as follows: 0.0-8.0 min from 45% B to 99% B, 8.0-9.0 min at 99% B, 9.0-9.1 min to 45% B, and 9.1-10.0 min at 45% B. Electrospray ionization was performed inLpositiveLor negativeLion mode.

We applied the following settings: curtain gas at 40 psi (negative mode) and curtain gas at 40 psi (positive mode); collision gas was set at 9; ion spray voltage at 2000 V (positive mode) or - 2000 V (negative mode); temperature at 250°C (positive mode) or 450°C (negative mode); ion source Gas 1 at 40 psi; ion source Gas 2 at 70 psi; entrance potential at 10 V (positive mode) or -10 V (negative mode); andL collision cell exit potential at 15 V (positive mode) or -15 V (negative mode). Data acquisition was performed in multiple reaction monitoring mode (MRM) with the collision energy (CE) values reported in **Supplementary Tables 1 and 2**. Area ratios of endogenous lipids and surrogate internal standards were quantified usingLSCIEX OS 3.1 (Sciex).

##### Metabolomics analysis

Metabolites analyses were performed by liquid chromatography on an ExionLC (Sciex) coupled with electrospray mass spectrometry TripleQuad 7500 (Sciex). For each analysis, 1 µL of the sample was injected on a Premier BEH amide 1.7 µm, 2.1×150 mm column (Waters) using a flow rate of 0.40 mL/min at 40°C. Mobile phase A consisted of water with 10 mM ammonium formate + 0.1% formic acid. Mobile phase B consisted of acetonitrile with 0.1% formic acid. The gradient was programmed as follows: 0.0–1.0 min at 95% B; 1.0–7.0 min to 50% B; 7.0–7.1 min to 95% B; and 7.1–10.0 min at 95% B. Electrospray ionization was performed in positive ion mode. We applied the following settings: curtain gas at 40 psi; collision gas was set at 9; ion spray voltage at 1600 V; the temperature at 350°C; ion source Gas 1 at 30 psi; ion source Gas 2 at 50 psi; entrance potential at 10 V; and collision cell exit potential at 10 V. Data acquisition was performed in MRM mode with the CE values reported in **Supp. Table 3**. Area ratios of endogenous metabolites and surrogate internal standards (**Supp. Table 3**) were quantified usingLSCIEX OS 3.1 (Sciex).

##### Analysis of glucosyl- and galactosyl-sphingolipids

Glucosyl- and galactosyl-sphingolipids analyses were performed by liquid chromatography ExionLC coupled to electrospray mass spectrometry TQ7500. For each analysis, 1 µL of sample was injected on a HALO HILIC 2.0 µm, 3.0 × 150 mm column (Advanced Materials Technology) using a flow rate of 0.48mL/min at 45°C. Mobile phase A consisted of 92.5/5/2.5 (vol/vol/vol) acetonitrile/isopropanol/water with 5 mM ammonium formate and 0.5% formic acid. Mobile phase B consisted of 92.5/5/2.5 (vol/vol/vol) acetonitrile/isopropanol/water with 5 mM ammonium formate and 0.5% formic acid. The gradient was programmed as follows: 0.0–2 min at 0% B, 2.1 min at 5% B, 4.5 min at 15% B, hold to 6.0 min at 15% B, up to 100% B at 6.1 min and hold to 7.0 min, drop back to 0% B at 7.1 min and hold to 8.5 min. Electrospray ionization was performed in positive ion mode. We applied the following settings: curtain gas at 40 psi; collision gas was set at 9 psi; ion spray voltage at 2250 V; temperature at 450°C; ion source Gas 1 at 40 psi; ion source Gas 2 at 70 psi; entrance potential at 10 V; and collision cell exit potential at 15 V. Area ratios of endogenous glucosyl- or galactosyl-sphingolipids and surrogate internal standards (Table 4) were quantified using SCIEX OS 3.1 (Sciex).

#### Immunohistochemistry

Paraformaldehyde fixed coronal tissue sections from each group of rAAV-injected mice were stained with the StrepTagII C23.21 antibody or cell type markers (neurons (NeuN), microglia (IBA-1), astrocytes (GFAP)) using previously published procedures. The full list of antibodies is listed in the key resources table. For this procedure, 40 µm coronal brain sections were processed using a free-floating method. For StrepTagII and CD68 antibodies, antigen retrieval with Citrate buffer pH 6.0 (30min) was performed for epitope retrieval. Sections were rinsed three times in phosphate-buffered saline containing 0.3% Triton-X100 (PBST) (0.1M Phosphate buffer, pH 7.4, 0.137 M NaCl, 0.3% Triton-X100) and reacted in PBST containing 1% hydrogen peroxide (30 min) to remove endogenous peroxidase activity, rinsed three times in PBST, blocked with 2.5% normal horse serum, and then incubated in optimal dilutions of antibody overnight with shaking at room temperature (RT). Sections were then rinsed three times, incubated in biotinylated anti-species immunoglobulin (Vector Laboratories) at 1:1000 for 2 hours at room temperature, rinsed three times and then incubated with avidin-biotin-peroxidase complex (ABC) (Vector Laboratories). Localization of bound antibody was visualized using avidin-biotin horseradish peroxidase (HRP) enzyme complex histochemistry and nickel ammonium sulfate-enhanced diaminobenzidine-HCl (100 µg/ml) (TCI Chemicals, Tokyo, Japan) as a substrate to produce a dark purple reaction product. Sections were then mounted on microscope slides and coverslipped with permanent mounting agent.

For detection of the twin-Strep tag on constructs, the StrepTagII C23.21 antibody was visualized using the Mouse on Mouse ImmPress HRP Polymer kit (Vector Laboratories) according to the manufacturer’s protocol. To Quantify IHC signal cortical, thalamic, and hippocampal regions were cropped from a whole coronal section image. Brain regions of interest analyzed using an automated pipeline created using CellProfiler (www.cellprofiler.org)^78^ for quantification.

#### Fluorescent immunohistochemistry

Double-color fluorescent immunohistochemistry was carried out to verify cellular co-localization of PGRN-expressing cells in hPGRN-*Grn^-/-^*mice with cellular antigenic markers such as neuronal marker (NeuN), microglial marker (Iba-1), and astrocyte marker (Gfap). Tissue sections were incubated with optimal dilutions of antibodies at 4 degrees overnight with shaking. After three washes (10 min each) in PBST, sections were incubated with optimal concentrations of fluorescent-labeled secondary antibodies. Bound primary antibodies were detected with Alexa Fluor 488-donkey anti-goat IgG, and Cy3-donkey anti-rabbit IgG. After three washes, sections were mounted, coverslipped with Immuno-mount fluorescent mounting media (Thermo Fisher) and imaged using Lecia DMi 8 microscope with a DFC9000 GT camera and system software (LAS X Life Science microscope software).

#### Cell Culture

HeLa *GRN^-/-^* cells were a gift from Dr. Shawn Ferguson (Yale) and generated using CRISPR as described.^79^ HeLa wild-type or *GRN^-/-^* cells were cultured in DMEM medium plus 10% fetal bovine serum (FBS) and 1% Pen/Strep and maintained at 37 °C with 5% CO2. 24 hours before collection DMEM media was replaced with OptiMEM media (Gibco).

#### HeLa Lysis and Media Collection

Cells were suspended in MES buffer (50mM MES pH6.5, 1% Triton, 150mM NaCl, 1XHALT PPI) 5uL for every 1mg cell pellet. Cells were then lysed on ice for 10 mins briefly vortexing every 3 minutes. Lysates were then spun at 600xg for 10 minutes and supernatant was collected. Conditioned media was collected from culture dish and spun at 500xg for 10 minutes to remove any cell debris.

#### Flash Frozen Mouse Brain Sample Processing for Immunoblot

To prepare samples for the immunoblot analysis of proteins, a novel protocol was developed in which approximately 40 mg of mouse hippocampal tissue from each sample was placed in a solution of PBS with added HALT phosphatase protease inhibitor (PPI) at a dilution of 1:2 (weight to volume). The PPI was diluted into the 1xPBS at 1:100. In the PBS + PPI solution, the sample was cut into smaller pieces with mini scissors. Once cut into smaller pieces, the sample is ready for further homogenization.

A bead lysis kit was used for the homogenization of these small, soft hippocampal samples. The samples were cut into pieces and still in the PBS + PPI solution were pipetted into 1.5 mL RINO screw-tap tubes (Next Advance) prefilled with zirconium oxide beads. Tubes were placed into the Bullet Blender (Next Advance) for homogenization.

Once homogenized, the solution was diluted 1:5 in RIPA lysis buffer supplemented with 1x HALT protease and phosphatase inhibitor. After 15 minutes, the solution was sonicated (30A; 2 seconds on; 8 seconds of rest; 10 seconds total sonication time/sample). After sonication, the solution was spun down in a centrifuge at 20,000xRCF at 4C for 10 minutes.

Protein concentration was measured with the bicinchoninic acid (BCA) assay, samples were frozen in aliquots at -80C.

#### Immunoblot

SDS/PAGE and immunoblotting of HeLa cell lysates, cell media, and mouse brain lysates were performed as described.^25, 80, 81^ Mouse brain running samples were prepared for immunoblot in 1X Laemmli loading buffer with 20 mM tris(2-carboxyethyl)phosphine (TCEP)) followed by denaturation at 70 degrees C for 15 minutes. For immunoblotting, protein samples were first separated on Bio-Rad TGX 4-20% 26-well gels at 100 V and transferred to a 0.2-micron nitrocellulose membrane using the Bio-Rad Trans-blot Turbo system. BulletBlock (Nacalai) for 30 minutes at room temperature membranes were incubated overnight at 4C with primary antibodies (STAR MATERIALS). Membranes were probed with anti-Histone H3 or anti-Beta tubulin antibodies and imaged on the Odyssey Fc (LI-COR), to normalize protein abundance between samples.

For hGRN2 and hGRN4 protein samples were separated using 4-20% BisTris gels run using MES buffer (Genscript) at 100V to resolve bands. Transfers were completed using Bio-Rad Trans-blot Turbo to a 0.2-micron nitrocellulose membrane, then blocked with Fish Serum Blocking Buffer (ThermoFisher) for 60 minutes at room temperature. Membranes were then incubated with primary antibodies (1ug/mL) overnight at 4C. All primary antibodies were diluted to a final concentration of 50% glycerol for long term storage at -20C. Near-infrared fluorescent secondary antibodies (diluted in TBST) or HRP-conjugated (diluted in 0.5% milk in TBST) antibodies (STAR Materials) were incubated for 1 hour at room temperature. For HRP visualization, blots were incubated in WesternSure PREMIUM Chemiluminescent Substrate (LI-COR) for 5 min before imaging. Near infrared or chemiluminescent blots were imaged using Odyssey Fc (LI-COR) and analyzed by Image Studio software 5.2 (LI-COR).

#### Alignment and Percent Identity

Granulin 1-7 amino acid sequences were accessed from Uniprot Human: P28799 (Table 5). Sequences were aligned using the msa R package with ClustalOmega.^82^ Alignments were visualized and consensus sequence calculated using ggmsa.^83^ Percent Identity of amino acids was calculated from the ClustalOmega hGRN alignment using Bio3D.^84–86^

| Granulin | UniProt Accession Number |
| --- | --- |
| hGRN1 | PRO_0000012695 |
| hGRN2 | PRO_0000012696 |
| hGRN3 | PRO_0000012697 |
| hGRN4 | PRO_0000012698 |
| hGRN5 | PRO_0000012699 |
| hGRN6 | PRO_0000012700 |
| hGRN7 | PRO_0000012701 |

#### Diagrams

All diagrams and representative figures were made using BioRender (biorender.com).

#### Statistics

##### Proteomics, lipidomics, and metabolomics

LC/MS data was Log2 transformed and ANOVAs for the following comparisons were performed (GFP-*Grn^-/-^* and GFP-*Grn+/+*) (GFP-*Grn^-/-^*and hPGRN-*Grn^-/-^)* (GFP-*Grn^-/-^* and hGRN2-*Grn^-/-^)* (GFP-*Grn^-/-^* and hGRN4-*Grn^-/-^*) p-values were adjusted using the Benjamini-Hochberg method. Abundance of individual proteins of interest were analyzed using One-Way ANOVA followed by Tukey’s post-hoc analysis. Variance was assessed using the Brown-Forsythe test, p=0.05 and the normality of GFP *Grn-/-* and GFP-*Grn+/+* samples was determined using the Shapiro-Wilk test p=0.05. PCA confidence intervals were analyzed using the PCAtools R package alpha set to 95%. The area ratios of endogenous lipids, metabolites, and surrogate internal standards were quantified using SCIEX OS 3.1.

Statistical analysis of significance for lipid and metabolite levels in samples was determined by One-way ANOVA with Tukey’s post-hoc analysis. * p < 0.05, ** p < 0.01, *** p < 0.001, **** p < 0.0001. (GFP-*Grn^-/-^* and GFP-*Grn+/+*) (GFP-*Grn^-/-^* and hPGRN-*Grn^-/-^)* (GFP-*Grn^-/-^* and hGRN2-*Grn^-/-^)* (GFP-*Grn^-/-^* and hGRN4-*Grn^-/-^*) comparisons are visualized in figures. <u>Immunohistochemistry and Lipofuscin</u>: IHC image quantification was performed single brain sections from 5 animals per group (N=5). Normality of GFP *Grn^-/-^* and GFP-*Grn^+/+^* samples was assessed using Shapiro-Wilk test p=0.05 and variance was assessed using Brown-Forsythe test p=0.05. Comparisons were conducted using Two-way ANOVA, one factor being brain region and the second being AAV treatment group. A full effect model was fitted and Tukey’s post-hoc analysis was completed comparing treatments groups to all other treatment groups within brain region.

##### Western Blot and ELISA Quantification

All blots were run using 5 individual animals per group (N=5) and normalized values were analyzed using One-Way ANOVA followed by Tukey’s post-hoc analysis. Variance and normality were assessed in the same manner as immunohistochemistry experiments. All regions were assessed independently. The hippocampal galectin-3 outlier was identified using Grubbs test p= 0.0001. ELISA data was analyzed using One-Way ANOVA followed by Tukey’s post-hoc test.

##### Visualization

All bar charts were produced in PRISM version 9 and heatmaps were made using Quickomics.^75^

## INCLUSION AND DIVERSITY

We support inclusive, diverse, and equitable research.

## SUPPLEMENTAL INFORMATION

Supplemental information can be found in attached PDFs.

