## Supplementary figures and images for "Granulins rescue inflammation, lysosome dysfunction, and neuropathology in a mouse model of progranulin deficiency"

### Supplemental Figures 1 through 4

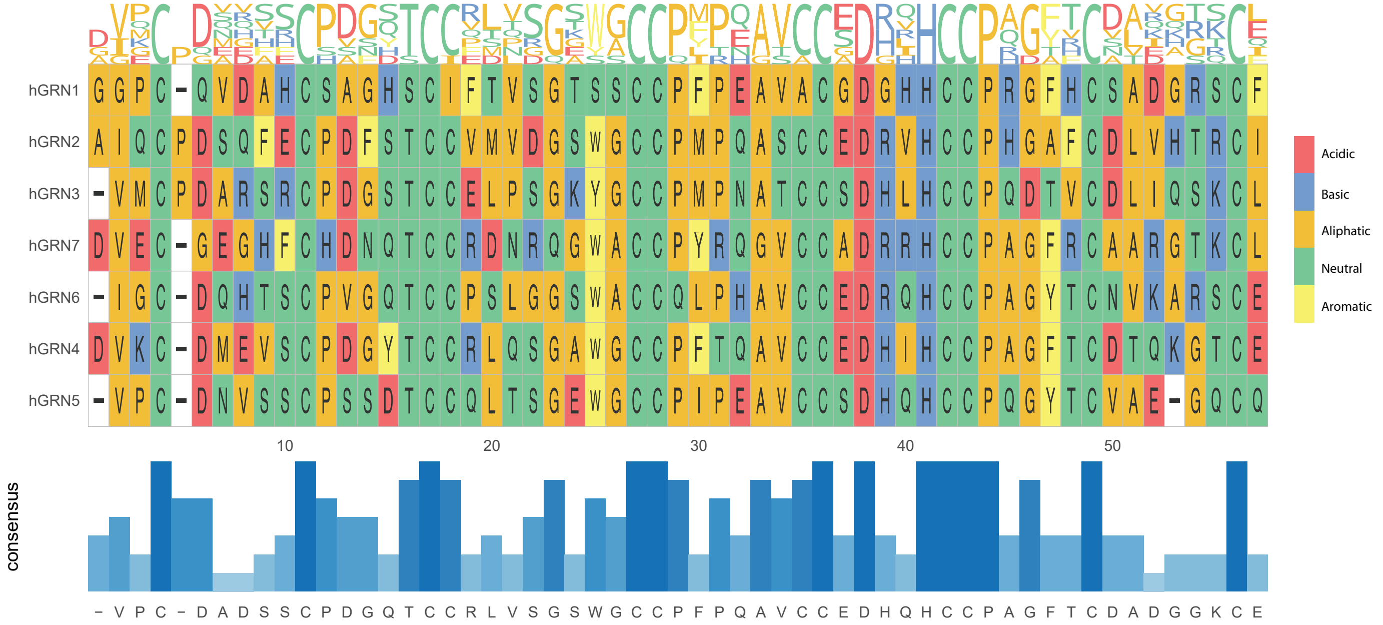

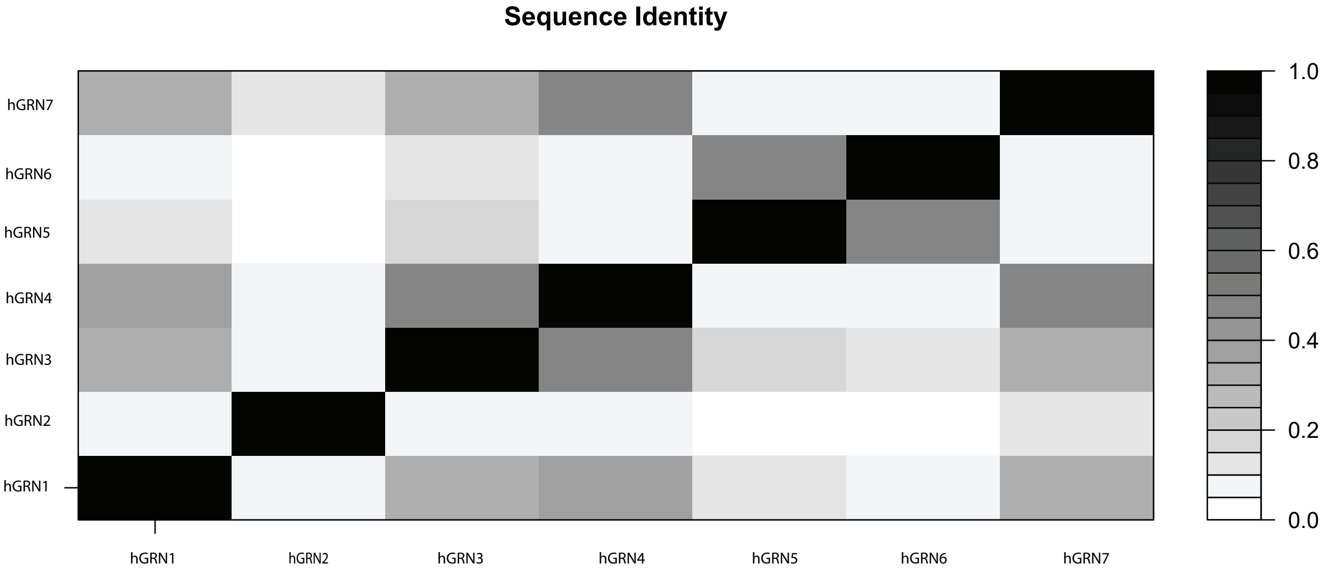

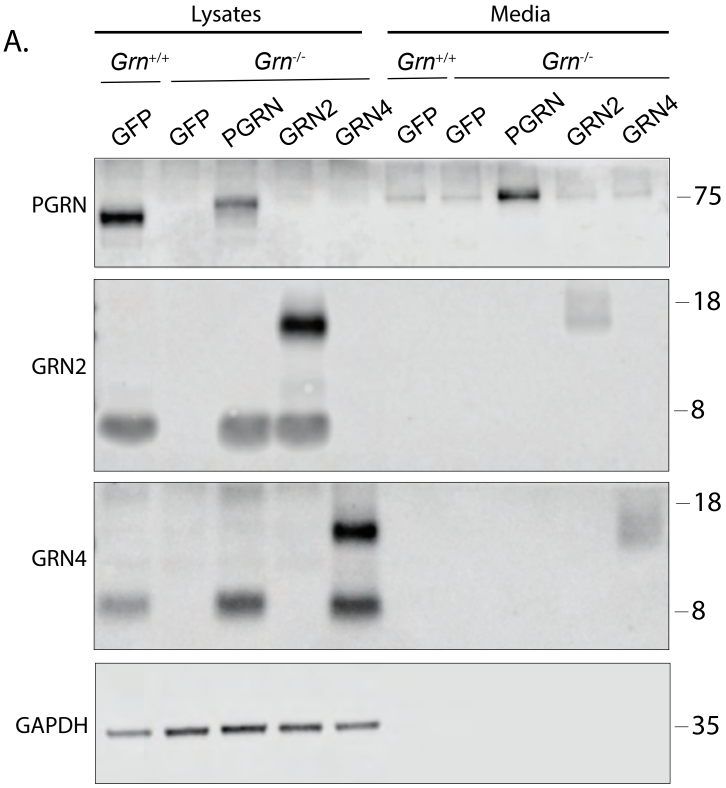

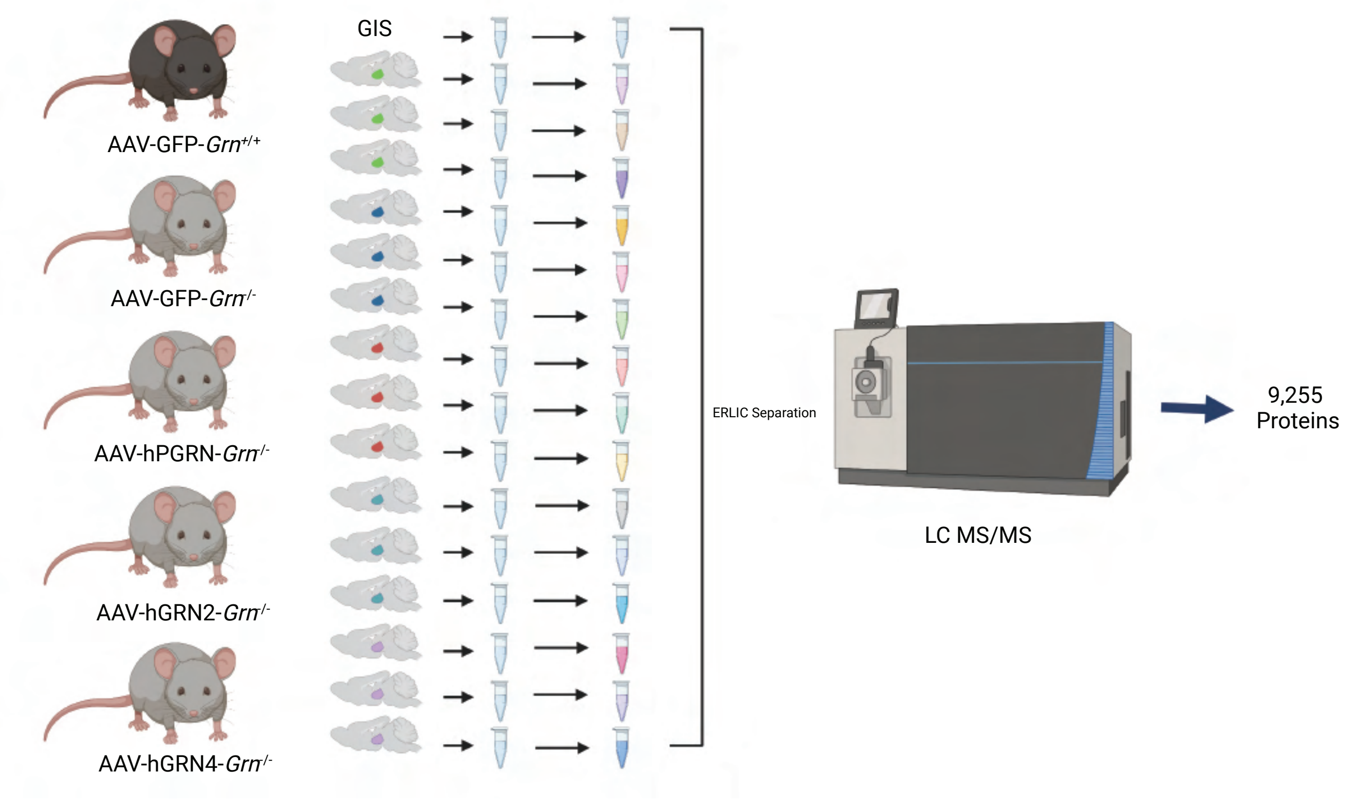

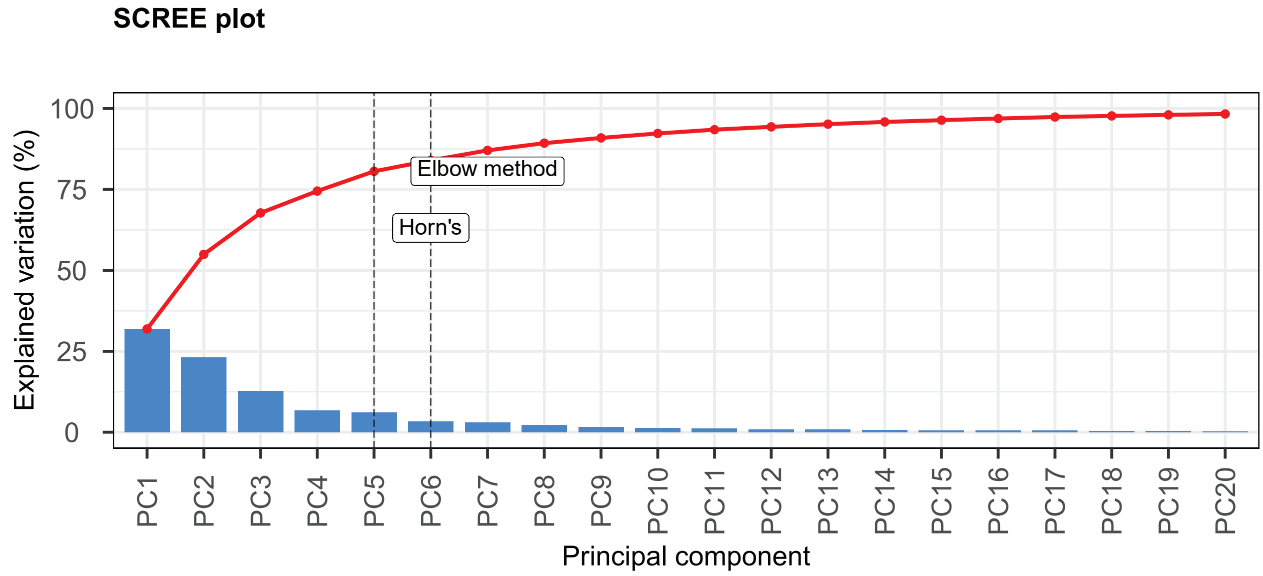

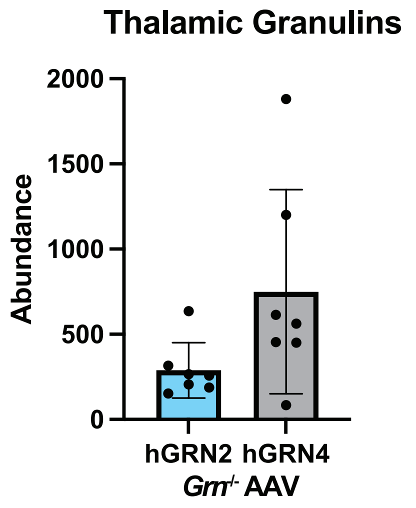

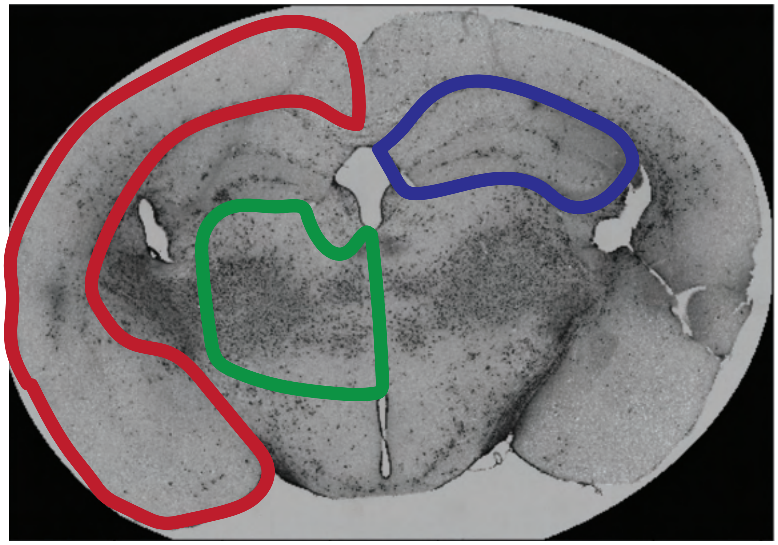

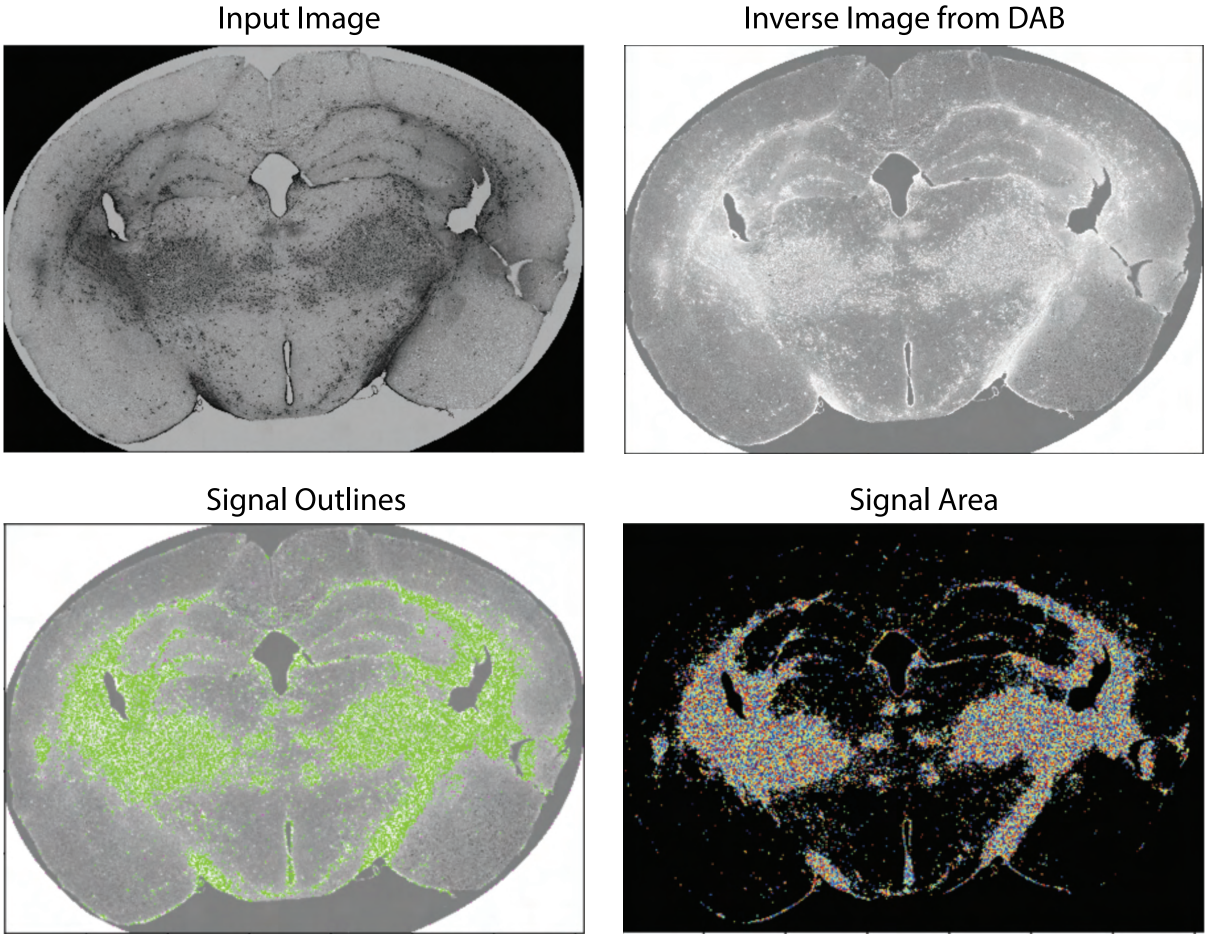
